# Microphase Separation Produces Interfacial Environment within Diblock Biomolecular Condensates

**DOI:** 10.1101/2023.03.30.534967

**Authors:** Andrew P. Latham, Longchen Zhu, Dina A. Sharon, Songtao Ye, Adam P. Willard, Xin Zhang, Bin Zhang

**Author notes:** These authors contributed equally to this work (Longchen Zhu, Dina A. Sharon).

## Abstract

The phase separation of intrinsically disordered proteins is emerging as an important mechanism for cellular organization. However, efforts to connect protein sequences to the physical properties of condensates, i.e., the molecular grammar, are hampered by a lack of effective approaches for probing high-resolution structural details. Using a combination of multiscale simulations and fluorescence lifetime imaging microscopy experiments, we systematically explored a series of systems consisting of diblock elastin-like polypeptides (ELP). The simulations succeeded in reproducing the variation of condensate stability upon amino acid substitution and revealed different microenvironments within a single condensate, which we verified with environmentally sensitive fluorophores. The interspersion of hydrophilic and hydrophobic residues and a lack of secondary structure formation result in an interfacial environment, which explains both the strong correlation between ELP condensate stability and interfacial hydrophobicity scales, as well as the prevalence of protein-water hydrogen bonds. Our study uncovers new mechanisms for condensate stability and organization that may be broadly applicable.

## INTRODUCTION

Biological condensates are found in both the cytosol^1–4^ and nucleus,^5–10^ playing essential roles in a variety of cellular processes^11,12^ from stress response^13^ to genome organization.^14,15^ Similar to membrane-bound organelles, they assemble a collection of molecules to raise the efficiency of sophisticated tasks. The lack of a membrane barrier allows fast material exchange between condensates and the cellular environment, rendering the molecular composition and stability of condensates more prone to regulations by external signals. ^16–18^

Intrinsically disordered proteins (IDPs) that promote multivalent, promiscuous interactions are key drivers of condensate formation. ^19–21^ Multiple mechanisms, including electrostatic, cation-*π*, *π*-*π*, hydrogen bonding, and hydrophobic interactions, contribute to the affinity among various chemical groups.^22–24^ Above a threshold concentration, as predicted by the Flory-Huggins theory, ^25^ interactions among IDPs can drive liquid-liquid phase separation (LLPS) to produce a highly concentrated phase that nevertheless remains dynamic. The simplicity of theory, while insightful, may prove insufficient for a comprehensive understanding of biological condensates.

Much remains to be learned regarding the connection between amino acid sequences and protein phase behaviors, or the so-called “molecular grammar” of protein condensates.^26–31^ These systems often exhibit complex viscoelastic behaviors and substructures with a layered organization,^32–35^ defying the mean-field assumption in the Flory-Huggins theory. Several more advanced theories have been introduced to better account for long-lived structural features that might persist in the polymer network. The sticker and spacer model^36,37^ has been adopted by Pappu and coworkers to account for strong and specific interactions that might form physical cross-links among protein molecules.^38^ In the meantime, the block copolymer theory may explain the microphase separation that can lead to layered structures.^39–45^ Because of their inherent assumptions, different theories are more appropriate for some systems but not others. High-resolution structural characterizations of condensates could offer further insight into their organizational principles and the applicable theories.

Further decoding the condensate grammar could also benefit from studies of simpler systems. Natural proteins often utilize highly complex amino acid sequences, rendering the attribution of an individual residue’s contribution to collective phenomena challenging. On the other hand, elastin-like polypeptides (ELPs), which are composed of pentapeptide repeats of valine-proline-glycine-X-glycine, serve as excellent models for studying protein phase behaviors. ELPs undergo phase separation upon heating with a lower critical solution temperature (LCST).^46–50^ Importantly, the guest residue X can be modified to any amino acid except proline with modern engineering approaches, ^51^ enabling systematic characterization of the impact on condensate stability and organization upon introducing specific residues.

We combine multiscale simulations with fluorescence lifetime imaging microscopy (FLIM) to probe condensates formed by diblock ELPs and investigate the contribution of amino acid composition to condensate stability. The simulation approach allows the sampling of largescale conformational rearrangements while providing atomistic resolution to quantify the solvation environment of individual residues. It succeeds in reproducing the stability of condensates, and reveals the formation of tubular network phases similar to gyroids. Such structures deviate from weak micelles that have been proposed for diblock ELPs in solution, and result in heterogeneous microenvironments within the condensate, which we verify *in-vitro* with FLIM. In addition, we find that the condensate stability exhibits a striking correlation with a hydrophobicity scale derived from interfacial transfer free energy, supporting an interfacial microenvironment of the condensate interior. The chemical specificity of the microenvironment is dictated by the peptide sequence that prevents a complete microphase separation between hydrophobic and hydrophilic groups. As a result, all condensates remain highly solvated after phase separation, producing water molecules that maintain hydrogen bonding with the exposed peptide backbone which persists even in residues with hydrophobic side chains. Our study takes a significant stride at connecting amino acid sequences with the structural and physical properties of biological condensates.

## RESULTS

### Multiscale simulations of diblock ELP condensates

Decoding the condensate grammar necessitates connecting protein sequences with collective physical properties. ELPs stand out because of their sequence simplicity and amenability for biological engineering, allowing systematic exploration and precise attribution of amino acid contributions. We focus on diblock ELPs of sequence (V-P-G-V-G)*_n_*-(V-P-G-X-G)*_n_*, which we abbreviate as V*_n_*X*_n_* (Fig. 1A). The guest residue in the first block is set as valine to promote phase separation, and we explore 20 systems in which every natural amino acid is substituted into the X position in the second block.

**Figure 1:**
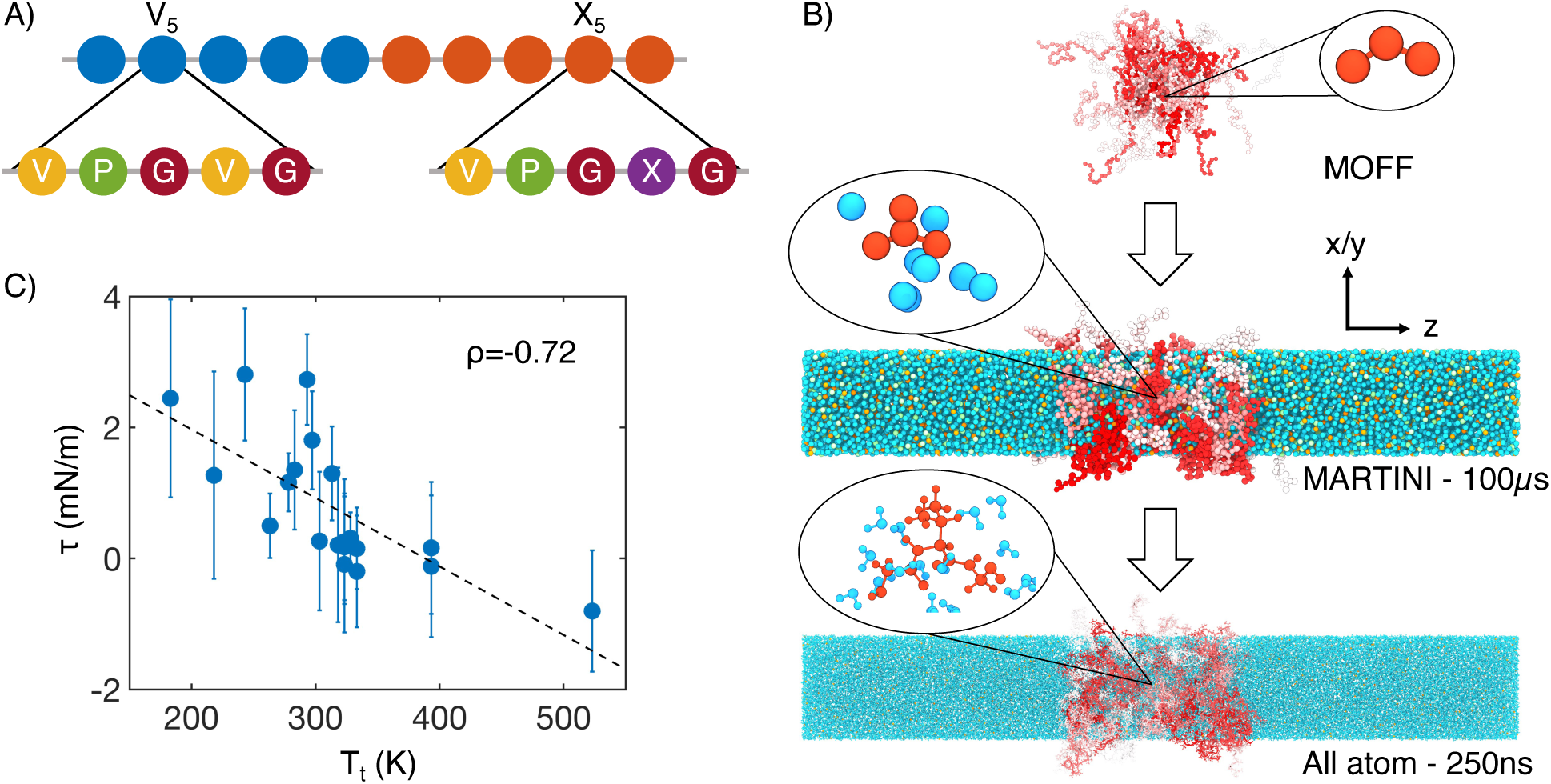
Multiscale simulations enable thermodynamic and structural characterization of ELP condensates. (A) Illustration of the sequence for the simulated diblock ELPs that consist of five-amino acid repeats, where X is substituted with a guest amino acid. (B) Overview of the three-step multiscale simulation approach that gradually increases the model resolution. Simulations of V_5_L_5_ were used to produce the example configurations at each step. Peptides are shaded red-white, while water molecules, chlorine, and sodium, are colored in blue, green, and orange, respectively. Inserts of a G-V-G repeat and surrounding water molecules are shown to indicate the resolution of each model. (C) Correlation between the simulated surface tension (*τ*) of 20 ELP condensates and the transition temperatures (*T*_t_) of related systems.^46^ *ρ* is the Pearson correlation coefficient between the two data sets, and the dashed line is the best fit between simulation and experimental data. Error bars represent the standard deviation of estimates from five independent time windows.

We adopt a multiscale approach to balance accuracy and efficiency for simulating ELP condensates (Fig. 1B). The first stage of this strategy is to simulate condensate formation using a coarse-grained force field, MOFF,^52–55^ with one bead per amino acid and implicit solvation. These simulations are highly efficient for system relaxation and promoting largescale conformational rearrangements that occur over slow timescales. We further converted these *α*-carbon based models to approximately four heavy atoms per bead resolution, and introduced explicit water and ions for simulations with the MARTINI force field.^56^ The latest version of this force field (MARTINI3) provides a more balanced set of parameters for protein-protein interactions and has been applied to study biological condensates.^57–60^ These simulations further relax the protein configurations and the partition of water and counterions in condensed and dilute phases. Finally, we carried out explicit solvent all-atom simulations for 250 ns to produce an accurate characterization of the chemical environment of the condensates.

Utilizing the simulated condensate conformations, we computed various quantities to benchmark against experimental measurements. While the critical temperature has been widely used as a measure for condensate stability, determining it computationally is expensive. As an alternative, we computed the surface tension, *τ*, using 100-*µ*s-long MARTINI simulations performed with the NP_N_AT ensemble.^61^ As detailed in the *Supplemental Theory* in the Supporting information, an inverse relationship is expected between *τ* and the critical temperature, *T_c_*, for systems exhibiting LCSTs. We further approximate *T_c_* with the transition temperatures (*T_t_*) of ELP sequences,^46^ which are the temperatures at which ELPs undergo an LCST transition at a specified solution condition. *T_t_* was shown to be linearly proportional to *T_C_*.^62,63^ As expected, a negative correlation can be readily seen between computed surface tension and experimental *T_t_* (Fig. 1C). This observed negative correlation between *T_t_* and *τ* supports the simulation approach’s accuracy in reproducing the sequence-dependent changes in ELP phase behavior.

We note that the experimental values were determined using different ELP sequences from those simulated here, and the relation between the surface tension and *T_c_* is likely nonlinear,^64^ both contributing to the imperfect agreement. In another study,^65^ we adopted the same multiscale approach to determine the dielectric constant for several ELP condensates. The simulated values agree well with those determined using FLIM experiments.

### Microphase separation of ELP condensates

Upon validating the accuracy of the simulation protocol for reproducing collective properties of ELP condensates, we examined their structural organization. Hassouneh et al. argued that, in dilute solutions, ELP diblocks self-assemble into the so-called weak spherical micelles with dense cores and almost unstretched coronas.^66^ At phase separation conditions, whether similar structural features are preserved in condensates remains unclear.

We found that most ELP condensates undergo a microphase separation. Representative configurations for four typical systems from MARTINI simulations are shown in the top panels of Fig. 2, with the V blocks in blue and X in red. Overall, the microphase separation manages to orient the more hydrophobic blocks towards the condensate interior as opposed to the interface. The relative density of guest blocks near the center of the condensate positively correlates with their hydrophobicity, as can be seen in both MARTINI (Fig. S1 in the Supporting Information) and all-atom (Fig. S2 in the Supporting Information) simulations.

**Figure 2:**
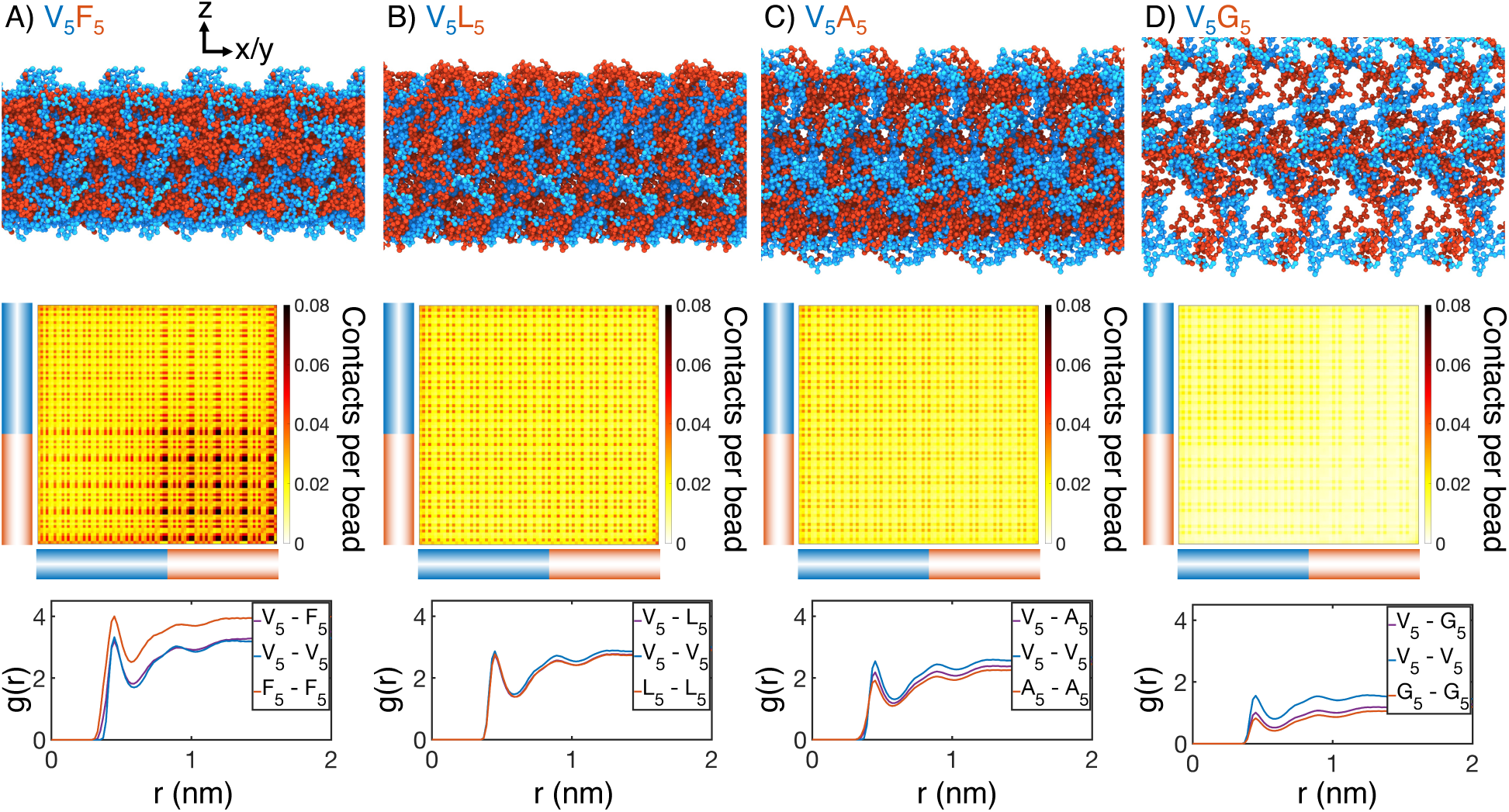
Internal organization of ELP condensates for (A) V_5_F_5_, (B) V_5_L_5_, (C) V_5_A_5_, and (D) V_5_G_5_. The uppermost panels present representative configurations from MARTINI simulations for each system. Periodic images along the x and y dimensions are shown for clarity, and the condensate-water interface is perpendicular to the z-axis. Only proteins are shown, with the X- and V-substituted halves of the peptides shown in red and blue, respectively. The central panels show contact maps between amino acids from different peptides. The blue and red bars indicate the V- and X-substituted half of the peptides. The lower panels plot the radial distribution functions, *g*(*r*), for amino acids only from the V-substituted half of the peptides (V_5_-V_5_), only from the X-substituted half of the peptides (X_5_-X_5_), and between the two halves (V_5_-X_5_). We limited the calculations to amino acid pairs from different peptides.

Surprisingly, microphase separation did not produce lamellar morphology as expected for block copolymers with equal volume fraction of the two blocks (Fig. S3 in the Supporting Information).^39–45,67–69^ In particular, the condensates appear to form gyroid-like structures, in which the V and X blocks form two interpenetrating networks (Fig. S4 in the Supporting Information). This morphology also differs from micelle-like structures seen in simplified hydrophobic-polar (HP) polymers.^70,71^ It promotes interfacial contacts while maintaining substantial self-interactions as well. Weak interfacial tension between different ELP blocks has also been noted by Hassouneh et al.^66^ These qualitative observations are insensitive to the polymer length and system size in our simulations (Fig. S5 in the Supporting Information). While longer diblock copolymers drive more prominent microphase separation, similar gyroid structures can also be observed. Notably, for more hydrophilic X blocks, the condensates begin to dissolve into a collection of micelle-like structures, consistent with the predictions by Hassouneh et al. in dilute solutions.^66^

Quantitative characterization of the condensate interior supports the presence of interpenetrating networks with substantial interfacial contacts as well. For example, significant contacts between V and X blocks can be readily seen in the inter-chain contact maps (Fig. 2, Fig. S6 in the Supporting Information). We further computed the radial distribution functions for amino acids in various blocks (Fig. 2, Fig. S7 in the Supporting Information). For all systems, we found that the probability of finding cross-block contacts V-X is often comparable to the intra-block contacts between X-X or V-V, depending on the relative hydrophobicity of V and X. For example, the more hydrophobic F blocks exhibit strong self-clustering, much more prominent than V-V blocks that are comparable to V-F blocks. On the other hand, clustering among V-V blocks and V-A blocks is more substantial than among A-A blocks. These results are again robust with respect to system setups (Fig. S5 in the Supporting Information).

To experimentally test the microphase separation behavior uncovered in simulations, we studied the micro-physicochemical properties of the V-end and X-end of the peptides. We constructed diblock peptides with the combination of 30 pentameric repeats of V block and X (A or G) block, namely V_30_A_30_ and V_30_G_30_ (*Experimental Sequences* Section in the Supporting Information). The amino-termini of V_30_A_30_ and V_30_G_30_ sequences were subsequently labeled with environmentally sensitive BODIPY or SBD fluorophores,^72,73^ whose lifetime could be measured to quantify the viscosity or polarity of the V-end (Fig. 3A, left panel).^74^ These probes have been reported to be only sensitive to single physicochemical properties.^65,73^ To avoid artifacts induced by fluorophore labeling, we usually used ELPs labeled with a low fraction of dyes. We also constructed A_30_V_30_ and G_30_V_30_ diblock peptides, wherein the viscosity or polarity of the A-end or the G-end could be measured by fluorophores that are attached at the amino-terminus (Fig. 3A, right panel).

**Figure 3:**
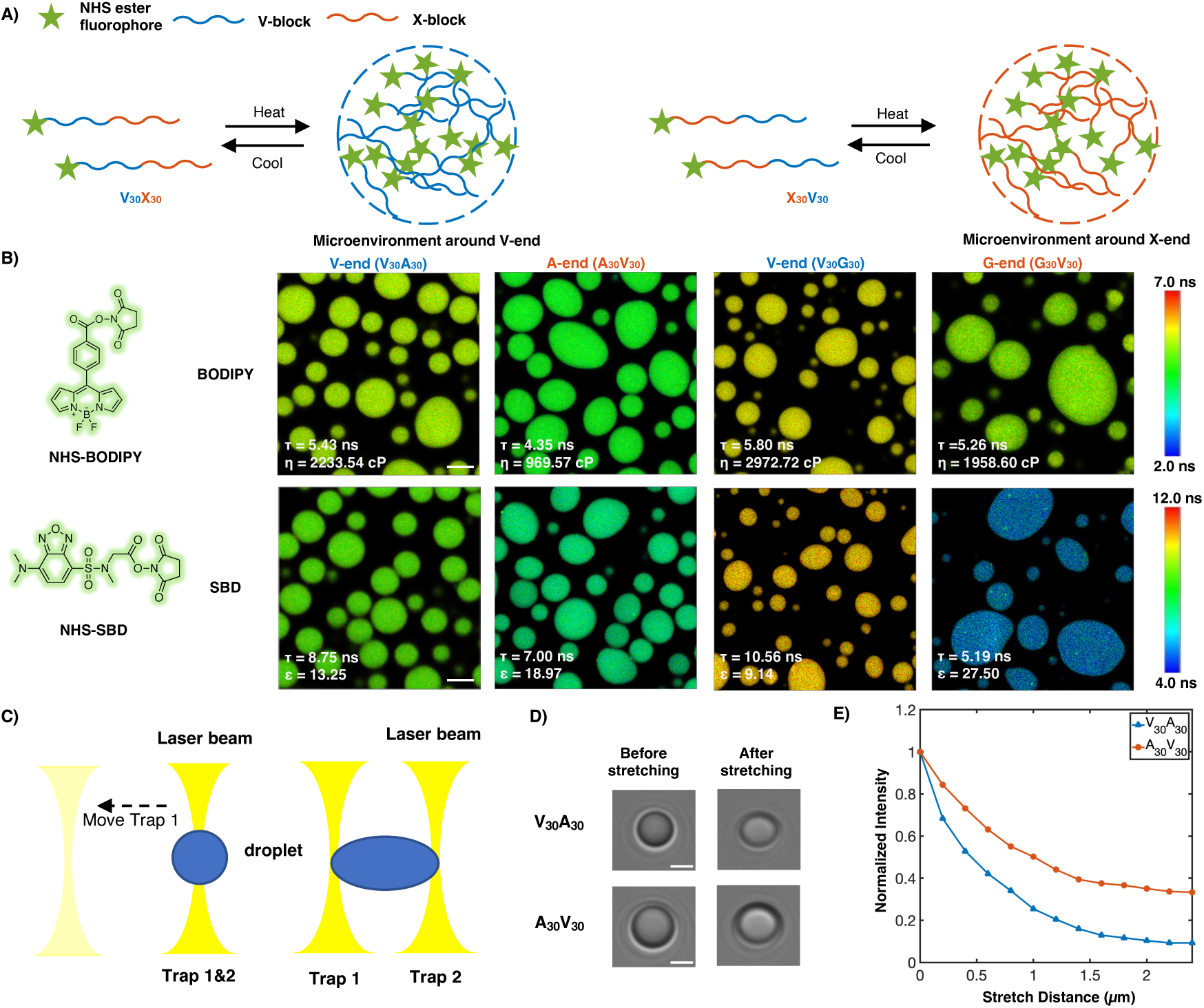
Experimental support of the microphase separation of ELP condensates. (A) Reversible ELP condensates formation via changing temperature. NHS ester fluorophores are attached at the amino-termini of V_30_X_30_ and X_30_V_30_ to detect the different micro-physicochemical properties between the V-end and the X-end. (B) Structures of NHSBODIPY and NHS-SBD, and FLIM images of V_30_A_30_, A_30_V_30_, V_30_G_30_ and G_30_V_30_ labeled with respective fluorophores. The fluorescence lifetime of each image is the average acquired from three independent experiments. Scale bar: 5 *µ*m. (C) Schematic diagram of optical tweezers stretching experiment. (D) Bright field images of V_30_A_30_ and A_30_V_30_ in the stretching experiment using the optical tweezer. Scale bar: 2 *µ*m. (E) Normalized droplets fluorescence intensity changes while stretching (red curve: V_30_A_30_; blue curve: A_30_V_30_). Fifteen droplets (at size *∼* 4*µ*m) were imaged and used for statistical analysis.

Using FLIM, we found that the lifetime of BODIPY for the V-end (5.43 ns) was longer than that for the A-end (4.35 ns), suggesting that the V-end indeed has a higher microviscosity than the A-end (*η_V_* = 2233.54 cp vs *η_A_*= 969.57 cp). Accordingly, the lifetime of SBD was longer for the V-end (8.75 ns) than the A-end (7.00 ns), indicating that the micropolarity of the V-end was lower than the A-end (*ɛ_V_* = 13.25 vs *ɛ_A_* = 18.97). These observations could be largely attributed to the greater extent of dehydration at the V-end due to its higher local peptide density. We further showed that the observed differences are not results of possible artifacts arising from any subtle distinctions between the two sequences V_30_A_30_ and A_30_V_30_ (*Experimental Characterization of ELP Condensates* Section in the Supporting Information, Fig. S8-S9 in the Supporting Information). Similar results were observed using the V-G sequences. FLIM experiments revealed that the V-end was more viscous than the G-end (*η_V_* = 2972.72 cp vs *η_G_*= 1958.60 cp) and the V-end was less polar than the G-end (*ɛ_V_* = 9.14 vs *ɛ_G_* = 27.50). These experimental observations provided the first line of evidence to support the microphase separation, as suggested by the simulation results. It is worth noting that the reported values, although related, may not quantitatively represent the steady-state viscosity. This discrepancy arises from the slow relaxation timescale inherent in ELP condensates with viscoelastic properties.

To further investigate these phenomena in real-time in motion, optical tweezers were used to stretch the droplet to observe the viscosity changes of different blocks. A single diblock condensate was captured by two optical traps (trap 1 and 2) at the same position, and it was stretched as trap 1 moved away from the original position (Fig. 3C-D). We used the peptides V_30_A_30_ and A_30_V_30_, with BODIPY labeled at their amino-terminus, to study the microviscosity changes where the fluorescence intensity decreases as the viscosity decrease. We found that the normalized fluorescence intensity of the V-end decreases greater than the A-end, indicating that the viscosity change of the V-end is larger (Fig. 3E). We anticipate that during stretching, condensates deform and molecules may deviate from the preferred chemical environment established with microphase separation. Such deviations impact the hydrophobic segments more, leading to more significant reduction in their interactions with surrounding molecules and correspondingly their viscosity.

### Frustration produces condensate interior with interfacial properties

The strong dependence of molecular organization on amino acid hydrophobicity suggests that the solvation environment of individual residues might be a determining factor for condensate stability. Indeed, as shown in the *Supplemental Theory* of the Supporting Information, the critical temperature is closely related to the free energy cost of transferring polymer beads from a solution state to a polymer-only environment. This transfer free energy is often used to quantify the hydrophobicity of amino acids.^75–81^ To explore their relationship more quantitatively, we compared the transition temperature for ELP condensates measured by Urry^46^ to several hydrophobicity scales. Fig. 4A shows that water-octanol and water-POPC-interface transfer free energies best correlate with the transition temperature. Meanwhile, other hydrophobicity measures, even including those derived specifically for disordered proteins, do not match as well. We note that our conclusion here is not dependent on the limited number of hydrophobicity measures examined in this study, and can be seen in a previous clustering of 98 different hydrophobicity scales.^82^

**Figure 4:**
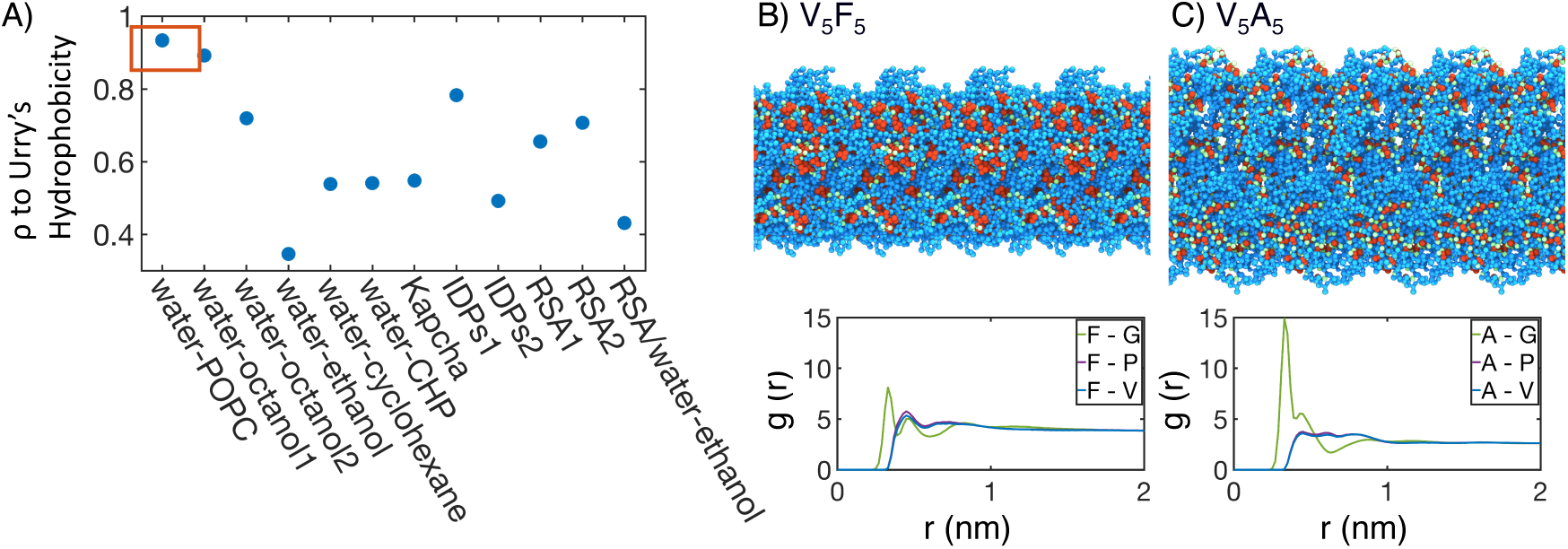
Interior of ELP condensates exhibits interfacial properties as a result of microphase separation. (A) Pearson correlation coefficient, *ρ*, between the stated measure of hydrophobicity and the condensate transition temperature, *T*_t_, measured by Urry.^46^ To remove discrepencies in which end of the scale is hydrophobic and which end is hydrophilic, all hydrophobicity scales, including the Urry scale, are first normalized such that 1 corresponds to the most hydrophobic residue and 0 corresponds to the least hydrophobic residue. Hydrophobicity measures considered include experimental measures of water-solvent transfer free energies (water-1-palmitoyl-2-oleoyl-sn-glycero-3-phosphocholine[POPC]-interface,^76^ water-octanol1,^77^ water-octanol2,^78^ water-ethanol,^78,79^ water-cyclohexane,^80^ and water-N-cyclohexyl-2-pyrrolidone[CHP]^81^), atomic level analysis of moieties within amino acids (Kapcha),^83^ computational approaches to estimate hydrophobicity within disordered proteins (IDPs1^84^ and IDPs2^85^), bioinformatics techniques that approximate the burial propensity, or relative solvent accessibility (RSA) of amino acids (RSA1^86^ and RSA2^87^), and a method that mixed protein burial fraction with water-vapor transfer free energy (RSA/water-vapor^88^). The red box highlights the high correlation between *T*_t_ and water-octanol or water-POPC-interface transfer free energies. (B, C) Condensate organization from MARTINI simulations for V_5_F_5_ (B) and V_5_A_5_ (C). The top panels provide protein-only views for simulated condensates, with the guest residue X, glycine adjacent to the guest residue, and the remaining residues colored red, green, and blue, respectively. Condensate images are repeated periodically in the x/y plane. The bottom panel shows the overall radial distribution function from the guest amino acid to those amino acids native to the ELP sequence.

The correlation analysis of the Urry transition temperature with various hydrophobicity scales supports a unique chemical environment emerging from the collective behaviors of condensates. To better understand the microenvironment surrounding guest residues in X blocks, we computed the radial distribution functions using MARTINI simulations. As shown in Fig. 4B-C and Fig. S10 in the Supporting Information, the guest residues are immediately surrounded with hydrophilic glycine residues as the nearest neighbors. In the second shell, more hydrophobic valine and proline residues appear. These profiles reveal the interfacial nature of the chemical environment surrounding the guest residues consisting of both hydrophobic and hydrophilic amino acids. This interfacial solvation environment is consistent with the strong correlation of the Urry scale with water-POPC-interface transfer free energy. Therefore, ELP condensates do not simply feature a hydrophobic interior that separates from hydrophilic residues as seen for folded proteins, and hence the poor correlation shown in Fig. 4A with the burial propensity of amino acids (RSA1, RSA2).

The ELP peptide sequences partly dictate the interfacial environment. For example, the guest residues are juxtaposed between two glycine residues in the pentamer (Fig. 1A), creating the immediate shell of hydrophilic residues. However, the presence of valine in the pentamer prevents the movement of GXG motifs to the interface due to the clustering of hydrophobic residues that prefer the condensate interior. Therefore, while microphase separation occurs in ELP condensates, frustration remains in the system. Hydrophilic residues cannot completely separate from hydrophobic ones due to constraints imposed by the amino acid sequence, creating unique microenvironments. Such a microenvironment arises from the collective behavior of many proteins and can deviate from individual chains,^89^ explaining the less-than-ideal correlation between the Urry scale and the hydrophobicity scales optimized to reproduce the conformational ensemble of IDP monomers.

### Solvation environment from atomistic simulations support interfacial properties

The interfacial environment uncovered in the previous section suggests the presence of significant solvation in the condensate interior. For condensates with lower surface tension, such as V_5_G_5_, the protein density near the center of the condensate is only around *∼* 30%. Even for the stable condensates with high surface tension, such as V_5_F_5_, the protein density near the center only reaches *∼* 70%. The relative density of protein, water, and ions in different systems generally follows the trend observed for the transition temperature (Fig. S11-S12 in the Supporting Information).

The large water mass fractions in the condensates prompt the question of the extent to which water is engaged in meeting the protein’s hydrogen bonding requirements. We determined the average number of hydrogen bonds per residue with ELP residues or water from extensive atomistic simulations. As shown in Fig. 5A, over 75% of the hydrogen bonds are between protein and water, in all of the condensates. Prior work on a different ELP condensate^47^ similarly indicated that water-protein hydrogen bonds constitute most of the hydrogen bonds found in the system. This finding highlights that ELPs, lacking noticeable secondary structures (Fig. S13 in the Supporting Information), are less capable of meeting its own hydrogen bonding needs, as compared to folded proteins.^90^ Therefore, high water concentration within the condensate is crucial to fulfilling the protein’s hydrogen bonding requirements.

**Figure 5:**
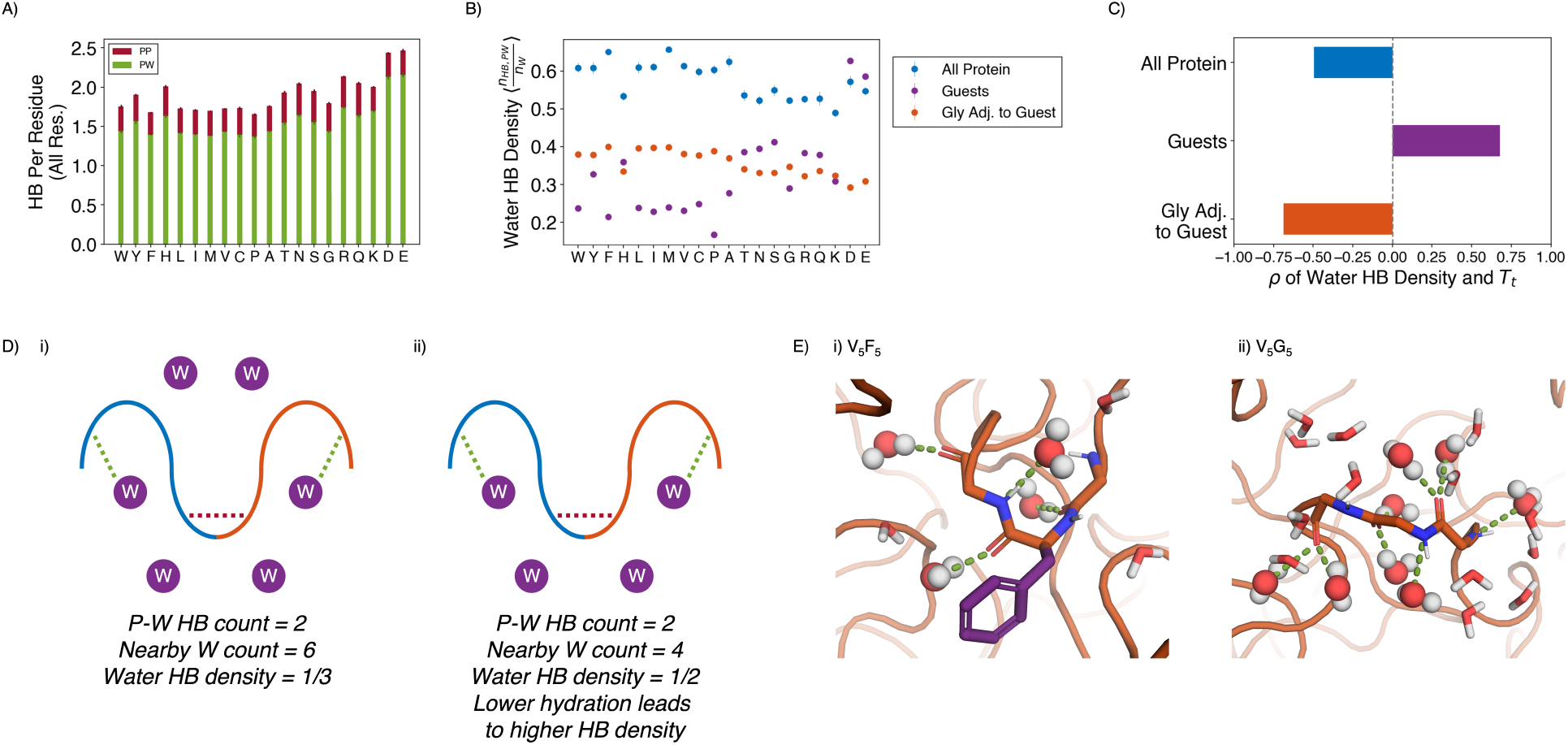
Water hydrogen bonding environment of condensates from all-atom simulations. (A) Bar chart depicting the average number of protein-water (PW, in green) and protein-protein (PP, in red) hydrogen bond (HB) per residue for each condensate system. The x-axis presents the guest residue of the system and is ordered by the Urry scale. Error bars represent standard deviations of four independent estimates. (B) Water HB density for each of the 20 condensates, for three different residue selections. The metric is shown for all residues in the system (blue), guest residues (purple), and glycine residues adjacent to the guest residue in the sequence (orange). The x-axis presents the guest residue of the system and is ordered by the Urry scale. Error bars represent standard deviations of four independent estimates. (C) Correlation coefficients of the water HB density shown in part B with Urry *T_t_*. (D) Schematics depicting two systems with low (i) and high (ii) water hydrogen bond density. The protein is depicted as blue and orange lines, and the water molecules are depicted as purple circles. Protein-protein and protein-water hydrogen bonds are drawn as red and green dashed lines respectively. (E) Visualizing hydrogen bond density with illustrative snapshots for V_5_F_5_ (i) and V_5_G_5_ (ii) condensate. Side chain carbon atoms are shown in purple. Water molecules near the protein are explicitly depicted, with oxygen and hydrogen atoms colored in red and white. Water molecules hydrogen bonded to the protein are depicted as spheres (with hydrogen bonds shown in green lines).

Strikingly, given the importance of water molecules, the average number of hydrogen bonds per residue remained relatively constant across condensates, even when the amount of water inside the condensate varied by almost two folds. Therefore, we hypothesize that each water molecule in more hydrophobic condensates must engage more in hydrogen bonding than each water in hydrophilic condensates. To test this hypothesis, we determined the water hydrogen bond density, defined as the ratio of the number of protein-water hydrogen bonds divided by the total number of water molecules near the condensate (Fig. 5D and 5E). As shown in Fig. 5B and 5C, the water hydrogen bond density indeed negatively correlates with the condensate transition temperature, *T_t_*, supporting our hypothesis that more hydrophobic condensates would have a higher hydrogen bond density.

The water hydrogen bond density also highlights an interfacial environment of blended hydrophobic and hydrophilic regions. For example, when we limit the analysis to water molecules near the guest residues, X, the hydrogen bond density now becomes positively correlated with *T_t_*. Therefore, locally, protein-water interactions are strongly influenced by the chemistry of side chains, and water molecules have fewer hydrogen bonding opportunities with more hydrophobic residues. However, the opposite trend is observed for the hydrogen bond density near glycine residues that immediately surround the guest residues. Therefore, the exposed backbones retain water molecules even when hydrophobic side chains do not provide hydrogen bonding opportunities, and the inseparability between the two further contributes to the observed interfacial property of condensates.

As a complementary measure of the solvation environment of individual residues, we computed the relative solvent accessibility (RSA) for the guest amino acids using atomistic simulations (Fig. 6A-B). The RSA measures the solvent-accessible surface area normalized by the maximum possible solvent exposure. RSA values in ELP condensates are much lower than those computed from all-atom simulations of ELP monomers, consistent with the increase of polymer density upon phase separation. However, they are almost always higher than the values estimated for folded proteins due to many water molecules roaming inside the condensates.^87^ Therefore, while ELP condensates provide a mechanism to shield hydrophobic amino acids from solvent, they are less effective in doing so than folded proteins.

**Figure 6:**
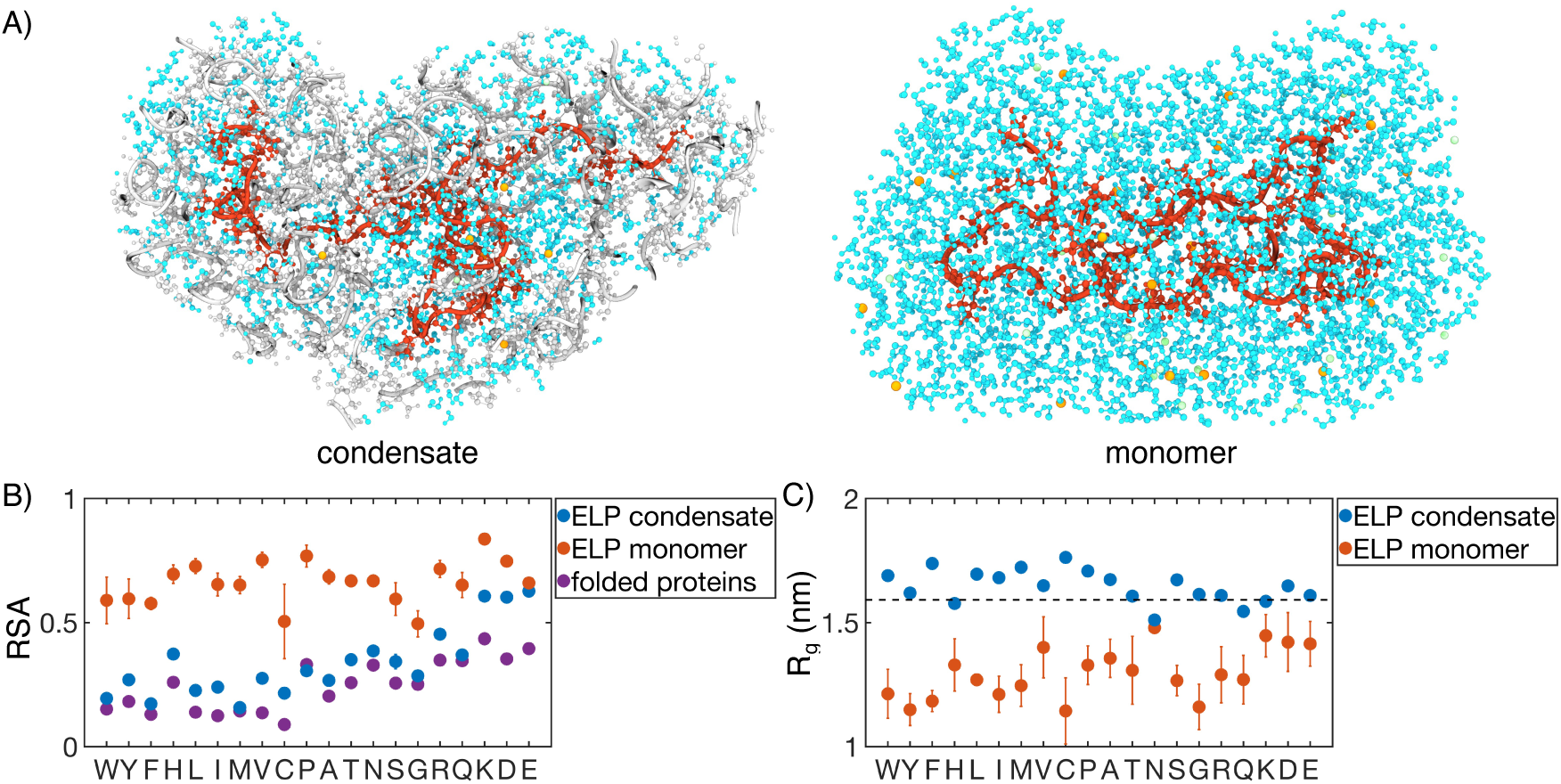
Comparison of protein conformation and solvation between ELP condensates and monomers. (A) Zoom in view of a single peptide chain (red) from atomistic simulations of V_5_A_5_ condensate (left) and monomer (right). Atoms within one nm of the peptide are shown, including water (cyan), chlorine (green), sodium (orange), and protein atoms from other chains (gray). (B) RSA of guest residues estimated from atomistic simulations of condensates (blue) and monomers (orange). For comparison, the corresponding values estimated using folded proteins are included as purple dots. ^87^ Error bars represent the standard deviation of four independent time estimates. (C) Comparison of the radius of gyration (*R_g_*) for peptides estimated from atomistic simulations of condensates (blue) and monomers (orange). The dashed line represents the expected *R_g_* of an ideal chain, utilizing values that have been previously suggested for IDPs.^91,92^ Error bars represent the standard deviation of four independent time estimates.

While the amino acid RSA values increase upon phase separation, unlike protein folding, the increase does not result from polymer collapse. We observe a significant expansion for almost all the studied peptides upon condensation, supporting the notion that water is a poor solvent for ELP (Fig. 6C). The expansion is most evident for more hydrophobic ELPs, the radii of gyration of which are now comparable to that of an ideal Gaussian chain. Notably, the sizes of the two separate peptide blocks are similar in most systems, supporting an interfacial environment that can accommodate both hydrophobic and hydrophilic residues (Fig. S14 in the Supporting Information).

## CONCLUSIONS and DISCUSSION

We carried out multiscale simulations to elucidate the connection between protein sequences and emergent properties of ELP condensates. The multiscale approach overcomes challenges in sampling slow conformational rearrangements to produce equilibrium atomistic condensate configurations. The simulations successfully reproduced condensate stability variation upon amino acid substitution. While our study is performed at set salt concentration and temperature to isolate the contributions of amino acid hydrophobicity to condensate organization, future studies may consider implementing temperature^50^ or salt^93^ dependent models to explore how solution conditions affect the organization of ELP condensates.

We found that diblock ELPs undergo microphase separation, with the more hydrophobic blocks localizing towards the condensate interior. The more hydrophobic blocks increase the local protein density, resulting in higher viscosity and lower dielectric constants, as determined in FLIM experiments. In contrast to the interior of folded proteins, these blocks are significantly solvated with water occupying over 30% of mass density. Unlike the weak micelles formed in solution,^66^ ELP condensates exhibit gyroid-like morphologies that promotes contacts between blocks. Further studies on the thermodynamic stability of these morphologies and comparing with predictions from the self-consistent field theory shall provide more insights into the driving forces for their emergence.

Our study uncovered a remarkably simple grammar for determining the stability of ELP condensates. A single quantity specific to the guest residue suffices to predict the emergent properties of the condensate, as evidenced by the strong correlation of the transition temperatures with the transfer free energies. The lack of higher-order effects is striking, supporting a mean field approximation to account for the contribution of individual residues in the system. However, the hydrophobicity scale is itself sensitive to the local microenvironment that is sequence specific. Such a microenvironment arises from the collective behavior of many proteins, can deviate from that of individual chains, and is likely sensitive to the solution conditions,^94^ which are held constant in our study. Future work on systems with double amino acid substitutions or changes to salt concentration or temperature could elucidate the generality of the mean field interpretation and the additivity of individual contributions.

While the transferability of the extracted rule for condensate stability to other systems remains to be shown, there are reasons to be hopeful. For instance, we anticipate that the microphase separation that produces the interfacial properties is a general principle for condensate organization. For most IDPs, the hydrophobic and hydrophilic residues are often interspersed throughout the chain to prevent the collapse of individual proteins^95,96^ and, correspondingly, the complete phase separation as in the lamellar phase. In addition, the hydrophilic backbones are expected to be exposed due to a lack of secondary structure formation, contributing to the interfacial property in the presence of hydrophobic side chains. As such, we anticipate that future research will discover the broad presence of interfacial environments within microphase separated motifs, as observed here.

## METHODS

### Molecular dynamics simulation details

To better understand the properties of biological condensates, we performed simulations on 40 diblock ELPs with the sequence (V-P-G-V-G)*_n_*-(V-P-G-X-G)*_n_*. We set *n* = 5 to balance computational efficiency and polymer topological effects. As shown in Fig. S5 in the Supporting Information, the results presented in the main text are qualitatively robust with respect to the system size and peptide length.

The multiscale simulations of ELP condensates started with an implicit solvent model, MOFF,^52^ and a similar homopolymer model, to generate initial configurations for each system. Next, we converted the *α−*carbon only configurations into those consistent with the MARTINI force field^57^ for coarse-grained, explicit solvent simulations. These simulations measured the coexistence between the condensate and solvent, lasted for 100 *µ*s, and were conducted in the NP_N_AT ensemble. Finally, we converted the MARTINI configurations into atomistic structures for simulations with CHARMM36m^97^ force field, with protein-water interactions shifted to better characterize IDPs. Atomistic simulations were conducted in the NP_N_AT ensemble and lasted 250 ns. All simulations were conducted using the software GROMACS^98^ and are described in detail in the *Condensate simulation details* section of the *Supporting Information*.

In addition to the condensates, we performed atomistic simulations of ELP monomers. These simulations began from RoseTTAFold predictions of the peptide structure ^99,100^ and lasted for 1 *µ*s in the NPT ensemble. Full details are available in the *Monomer simulation details* section of the *Supporting Information*.

### Analysis of condensate simulations

#### Surface tension

In explicit solvent simulations, two coexisting phases are present, in which proteins form a dense slab in the x-y plane, and the solvent creates a dilute phase in the z-dimension.^101,102^ We controlled the pressure normal to the protein-solvent interface by performing simulations in the NP_N_AT ensemble.^61^ The surface tension (*τ*) was calculated from these simulations as

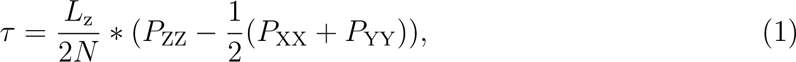

where *L*_z_ is the length of the Z-dimension and *N* is the number of protein droplets. *P*_XX_, *P*_YY_, and *P*_ZZ_ are the components of the pressure tensor along the three axes.

#### Clustering

Since the condensate can diffuse along the z-axis, i.e., the direction perpendicular to the interface, we aligned the simulated configurations using the position of the largest cluster of peptide molecules. To find the largest cluster, we first computed a center of mass contact matrix between peptide molecules with a distance cutoff of 4 nm. Then, using a depth first search algorithm,^103^ we identified the size and location of the largest protein cluster. The system was then shifted such that the z coordinate of the center of mass of the largest cluster is at zero. We implemented the alignment using the software MDAnalysis.^104,105^

#### Contact map

Protein contact maps were calculated using a cutoff distance of 0.6 nm. Specifically, two residues were defined as in contact if any pair of their coarse-grained beads were within 0.6 nm.

#### Radial distribution function

Radial distribution functions were calculated using the InterRDF function of MDAnalysis between 0.00001 nm and 2 nm. For analyzing the microphase separation of each condensate, we divided each protein into the V-substituted blocks and X-substituted blocks. This radial distribution function excluded intra-chain contacts to better capture how inter-chain contacts determine condensate organization. To analyze the origin of chain frustration within diblock phase separation, we computed the radial distribution function of each guest amino acid to the amino acids native to the sequence. Intra-chain contacts were included to fully capture the local environment experienced by the guest amino acid.

#### Relative solvent accessibility

The relative solvent accessibility (RSA) was measured based on the solvent accessible surface area (SASA) of guest amino acids determined from all atom simulations. We first calculated the SASA for each residue using GROMACS tools sasa, with a probe radius of 0.14 nm.^106^ We then averaged across all repeats of guest amino acids, and normalized the SASA by the maximum possible value for a given amino acid to compute RSA^87^ (see Table S1).

#### Radius of gyration

The estimated *R_g_* from atomistic simulations were compared to the following analytical expression^91^

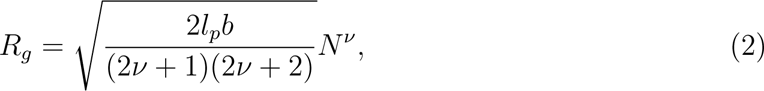

where *N* is the length of the polymer, *b* = 0.38nm is the bond length, and *l_p_* = 0.40nm is the persistence length.

#### Secondary structure

Secondary structure for protein condensates was calculated from atomistic simulations using the GROMACS do dssp command.^107,108^ Fig. S13 in the Supporting Information displays the ordered fraction of secondary structures, which includes *α*-helices, *β*-sheets, *β*-bridges, and turns.

#### Hydrogen bonding

Protein-protein and protein-water hydrogen bonds were determined using the hydrogenbonds package within MDAnalysis.^104,105,109^ A hydrogen bond was defined as having a donor heavy atom within 3.30 Å of an acceptor heavy atom and a donor heavy atom-hydrogen atom-acceptor heavy atom angle of 135.0*^◦^* or larger. The nearby water count for each analysis selection was determined by counting the water molecules whose oxygen atoms were within 4.0 Å of the heavy atoms of this selection. Water hydrogen bond density for each residue selection for each frame was computed as the ratio of protein-water hydrogen bonds to the number of nearby water molecules (Fig. S15 in the Supporting Information).

### Experimental methods

#### Fluorescence lifetime imaging microscopy (FLIM)

The proteins of interest were labeled with NHS ester fluorophore. We used ELPs with 1% BODIPY labels or 2.5% SBD labels to form condensates, which avoid the artifacts induced by fluorophores. Droplets were formed with the final concentration of 70 *µ*M ELP in 2 M NaCl for V-A and 1.5 M NH4SO4 for V-G diblock, respectively. A drop of droplets containing solution was placed on a 0.17 mm coverslip with a 500 *µ*m spacer. Images were acquired by Leica Falcon Fluorescence Microscope equipped with Wil pulse laser and 63X/0.12 oil-immersion objective. The BODIPY was excited at 488 nm and the SBD was excited at 448 nm. The fluorescence lifetime fitting and image analysis were performed in LAS X and Image J.

#### Optical tweezer experiments

The protocol describes the preparation and manipulation of a sample containing a BODIPY-labeled ELP protein using an optical tweezer, specifically the C-Trap from LULICKS. The goal is to form a single large condensate from small droplets and then stretch it using the optical tweezers while monitoring its fluorescence intensity. The first step is to mix the BODIPY-labeled ELP protein with NaCl solution to obtain a final concentration of 20 *µ*M protein in 2 M NaCl. The sample is then loaded into the flow channel of the optical tweezer. The dual trap mode of the C-Trap is then used to move the two traps to the same position. The sample is flushed until a single large condensate of about 4 *µ*m is caught by the dual trap via the fusion of small droplets. The traps are then moved along with the droplet to another flow channel containing 2 M NaCl buffer. The channel and pressure are turned off to avoid any additional force on the droplet. The next step is to stretch the droplet by moving trap 1 0.2 *µ*m each time. Fluorescent images of each stretch are taken using a 488 nm excitation laser. The mean fluorescence intensity of the whole droplet is then quantified using Image J.

## Acknowledgement

This work was supported by the National Institutes of Health (Grant R35GM133580, B.Z.), the Research Center for Industries of the Future (RCIF) at Westlake University (X.Z), PEW Biomedical Scholars Program 00033066 (X.Z.), and the National Science Foundation under CHE-1654415 (A.P.W.). A.L. acknowledges support from the National Science Foundation Graduate Research Fellowship Program (grant number 1745302). D.S. acknowledges support from the Hertz Foundation and the National Science Foundation Graduate Research Fellowship Program (grant number 2141064).

## Supporting Information Available

The Supporting Information is available as a separate file for download.

## References

(1) Brangwynne, C. P.; Eckmann, C. R.; Courson, D. S.; Rybarska, A.; Hoege, C.; Gharakhani, J.; Julicher, F.; Hyman, A. A. Germline P Granules Are Liquid Droplets That Localize by Controlled Dissolution/Condensation. Science 2009, 324, 1729–1732.

(2) Folkmann, A. W.; Putnam, A.; Lee, C. F.; Seydoux, G. Regulation of biomolecular condensates by interfacial protein clusters. Science 2021, 373, 1218–1224.

(3) Patel, A. et al. A Liquid-to-Solid Phase Transition of the ALS Protein FUS Accelerated by Disease Mutation. Cell 2015, 162, 1066–1077.

(4) Ma, W.; Mayr, C. A Membraneless Organelle Associated with the Endoplasmic Reticulum Enables 3’UTR-Mediated Protein-Protein Interactions. Cell 2018, 175, 1492– 1506.e19.

(5) Feric, M.; Vaidya, N.; Harmon, T. S.; Mitrea, D. M.; Zhu, L.; Richardson, T. M.; Kriwacki, R. W.; Pappu, R. V.; Brangwynne, C. P. Coexisting Liquid Phases Underlie Nucleolar Subcompartments. Cell 2016, 165, 1686–1697.

(6) Larson, A. G.; Elnatan, D.; Keenen, M. M.; Trnka, M. J.; Johnston, J. B.; Burlingame, A. L.; Agard, D. A.; Redding, S.; Narlikar, G. J. Liquid droplet formation by HP1*α* suggests a role for phase separation in heterochromatin. Nature 2017, 547, 236–240.

(7) Strom, A. R.; Emelyanov, A. V.; Mir, M.; Fyodorov, D. V.; Darzacq, X.; Karpen, G. H. Phase separation drives heterochromatin domain formation. Nature 2017, 547, 241– 245.

(8) Sabari, B. R. et al. Coactivator condensation at super-enhancers links phase separation and gene control. Science 2018, 361, eaar3958.

(9) Leicher, R.; Osunsade, A.; Chua, G. N. L.; Faulkner, S. C.; Latham, A. P.; Watters, J. W.; Nguyen, T.; Beckwitt, E. C.; Christodoulou-rubalcava, S.; Young, P. G.; Zhang, B.; David, Y.; Liu, S. Single-stranded nucleic acid binding and coacervation by linker histone H1. Nat. Struct. Mol. Biol. 2022, 29, 463–471.

(10) Latham, A. P.; Zhang, B. On the stability and layered organization of protein-DNA condensates. Biophys. J. 2022, 121, 1727–1737.

(11) Banani, S. F.; Lee, H. O.; Hyman, A. A.; Rosen, M. K. Biomolecular condensates: Organizers of cellular biochemistry. Nat. Rev. Mol. Cell Biol. 2017, 18, 285–298.

(12) Uversky, V. N. Intrinsically disordered proteins in overcrowded milieu: Membrane-less organelles, phase separation, and intrinsic disorder. Curr. Opin. Struct. Biol. 2017, 44, 18–30.

(13) Riback, J. A.; Katanski, C. D.; Kear-Scott, J. L.; Pilipenko, E. V.; Rojek, A. E.; Sosnick, T. R.; Drummond, D. A. Stress-Triggered Phase Separation Is an Adaptive, Evolutionarily Tuned Response. Cell 2017, 168, 1028–1040.e19.

(14) Lin, X.; Qi, Y.; Latham, A. P.; Zhang, B. Multiscale Modeling of Genome Organization with Maximum Entropy Optimization. J. Chem. Phys. 2021, 155, 010901.

(15) Sabari, B. R.; Dall’Agnese, A.; Young, R. A. Biomolecular Condensates in the Nucleus. Trends Biochem. Sci. 2020, 961–977.

(16) Hnisz, D.; Shrinivas, K.; Young, R. A.; Chakraborty, A. K.; Sharp, P. A. A phase separation model predicts key features of transcriptional control. Cell 2017, 169, 13– 23.

(17) Riback, J. A.; Zhu, L.; Ferrolino, M. C.; Tolbert, M.; Mitrea, D. M.; Sanders, D. W.; Wei, M. T.; Kriwacki, R. W.; Brangwynne, C. P. Composition-dependent thermodynamics of intracellular phase separation. Nature 2020, 581, 209–214.

(18) Klein, I. A. et al. Partitioning of cancer therapeutics in nuclear condensates. Science 2020, 368, 1386–1392.

(19) Hyman, A. A.; Weber, C. A.; Jülicher, F. Liquid-Liquid Phase Separation in Biology. Annu. Rev. Cell Dev. Biol. 2014, 30, 39–58.

(20) Boeynaems, S.; Alberti, S.; Fawzi, N. L.; Mittag, T.; Polymenidou, M.; Rousseau, F.; Schymkowitz, J.; Shorter, J.; Wolozin, B.; Van Den Bosch, L.; Tompa, P.; Fuxreiter, M. Protein Phase Separation: A New Phase in Cell Biology. Trends Cell Biol. 2018, 28, 420–435.

(21) Abyzov, A.; Blackledge, M.; Zweckstetter, M. Conformational Dynamics of Intrinsically Disordered Proteins Regulate Biomolecular Condensate Chemistry. Chem. Rev. 2022, 122, 6719–6748.

(22) Dignon, G. L.; Best, R. B.; Mittal, J. Biomolecular Phase Separation: From Molecular Driving Forces to Macroscopic Properties. Annu. Rev. Phys. Chem. 2020, 71, 1–23.

(23) Das, S.; Lin, Y. H.; Vernon, R. M.; Forman-Kay, J. D.; Chan, H. S. Comparative roles of charge, *π*, and hydrophobic interactions in sequence-dependent phase separation of intrinsically disordered proteins. Proc. Natl. Acad. Sci. U.S.A. 2020, 117, 28795– 28805.

(24) Murthy, A. C.; Tang, W. S.; Jovic, N.; Janke, A. M.; Seo, D. H.; Perdikari, T. M.; Mittal, J.; Fawzi, N. L. Molecular interactions contributing to FUS SYGQ LC-RGG phase separation and co-partitioning with RNA polymerase II heptads. Nat. Struct. Mol. Biol. 2021, 28, 923–935.

(25) Flory, P. J. Thermodynamics of high polymer solutions. J. Chem. Phys. 1942, 10, 51.

(26) Schuster, B. S.; Regy, R. M.; Dolan, E. M.; Kanchi Ranganath, A.; Jovic, N.; Khare, S. D.; Shi, Z.; Mittal, J. Biomolecular Condensates: Sequence Determinants of Phase Separation, Microstructural Organization, Enzymatic Activity, and Material Properties. J. Phys. Chem. B 2021, 125, 3441–3451.

(27) Choi, J. M.; Holehouse, A. S.; Pappu, R. V. Physical Principles Underlying the Complex Biology of Intracellular Phase Transitions. Annu. Rev. Biophys. 2020, 49, 107– 133.

(28) Wang, J.; Choi, J. M.; Holehouse, A. S.; Lee, H. O.; Zhang, X.; Jahnel, M.; Maharana, S.; Lemaitre, R.; Pozniakovsky, A.; Drechsel, D.; Poser, I.; Pappu, R. V.; Alberti, S.; Hyman, A. A. A Molecular Grammar Governing the Driving Forces for Phase Separation of Prion-like RNA Binding Proteins. Cell 2018, 174, 688–699.e16.

(29) Schneider, R.; Blackledge, M.; Jensen, M. R. Elucidating binding mechanisms and dynamics of intrinsically disordered protein complexes using NMR spectroscopy. Curr. Opin. Struct. Biol. 2019, 54, 10–18.

(30) Elbaum-Garfinkle, S.; Kim, Y.; Szczepaniak, K.; Chen, C. C.-H.; Eckmann, C. R.; Myong, S.; Brangwynne, C. P. The disordered P granule protein LAF-1 drives phase separation into droplets with tunable viscosity and dynamics. Proc. Natl. Acad. Sci. U.S.A. 2015, 112, 7189–7194.

(31) Kilgore, H. R.; Young, R. A. Learning the chemical grammar of biomolecular condensates. Nat. Chem. Biol. 2022,

(32) Latham, A. P.; Zhang, B. Molecular Determinants for the Layering and Coarsening of Biological Condensates. Aggregate 2022, e306.

(33) Mittag, T.; Pappu, R. V. A conceptual framework for understanding phase separation and addressing open questions and challenges. Mol. Cell 2022, 82, 2201–2214.

(34) Majumdar, A.; Krainer, G. Phase-separating RNA-binding proteins form heterogeneous distributions of clusters in subsaturated solutions. Proc. Natl. Acad. Sci. U.S.A. 2022, 119, e2202222119.

(35) Wu, T.; King, M. R.; Farag, M.; Pappu, R. V.; Lew, M. D. Single fluorogen imaging reveals spatial inhomogeneities within biomolecular condensates. bioRxiv 2023, 525727.

(36) Tanaka, F. Theory of Thermoreversible Gelation. Macromolecules 1989, 22, 1988–1994.

(37) Semenov, A. N.; Rubinstein, M. Thermoreversible Gelation in Solutions of Associative Polymers. 1. Statics. Macromolecules 1998, 31, 1373–1385.

(38) Harmon, T. S.; Holehouse, A. S.; Rosen, M. K.; Pappu, R. V. Intrinsically disordered linkers determine the interplay between phase separation and gelation in multivalent proteins. Elife 2017, 6, 1–31.

(39) Leibler, L. Theory of Microphase Separation in Block Copolymers. Macromolecules 1980, 13, 1602–1617.

(40) Bates, F. S.; Fredrickson, G. H. Block Copolymer Thermodynamics: Theory and Experiment. Annu. Rev. Phys. Chem. 1990, 41, 525–557.

(41) Matsen, M. W.; Schick, M. Stable and unstable phases of a diblock copolymer melt. Phys. Rev. Lett. 1994, 72, 2660–2663.

(42) Matsen, M. W.; Bates, F. S. Unifying weak- and strong-segregation block copolymer theories. Macromolecules 1996, 29, 1091–1098.

(43) Grason, G. M. The packing of soft materials: Molecular asymmetry, geometric frustration and optimal lattices in block copolymer melts. Phys. Rep. 2006, 433, 1–64.

(44) Swann, J. M.; Topham, P. D. Design and application of nanoscale actuators using block-copolymers. Polymers 2010, 2, 454–469.

(45) Shi, A. C. Frustration in block copolymer assemblies. J. Phys. Condens. Matter 2021, 33.

(46) Urry, D. W. Physical chemistry of biological free energy transduction as demonstrated by elastic protein-based polymers. J. Phys. Chem. B 1997, 101, 11007–11028.

(47) Rauscher, S.; Pomés, R. The liquid structure of elastin. eLife 2017, 6, e26526.

(48) Baul, U.; Bley, M.; Dzubiella, J. Thermal Compaction of Disordered and Elastin-like Polypeptides: A Temperature-Dependent, Sequence-Specific Coarse-Grained Simulation Model. Biomacromolecules 2020, 21, 3523–3538.

(49) Cinar, H.; Fetahaj, Z.; Cinar, S.; Vernon, R. M.; Chan, H. S.; Winter, R. H. Temperature, Hydrostatic Pressure, and Osmolyte Effects on Liquid–Liquid Phase Separation in Protein Condensates: Physical Chemistry and Biological Implications. Chem. Eur. J. 2019, 25, 13049–13069.

(50) Dignon, G. L.; Zheng, W.; Kim, Y. C.; Mittal, J. Temperature-Controlled Liquid–Liquid Phase Separation of Disordered Proteins. ACS Cent. Sci. 2019, 5, acscentsci.9b00102.

(51) McDaniel, J. R.; MacKay, J. A.; Quiroz, F. G.; Chilkoti, A. Recursive directional ligation by plasmid reconstruction allows rapid and seamless cloning of oligomeric genes. Biomacromolecules 2010, 11, 944–952.

(52) Latham, A. P.; Zhang, B. Consistent Force Field Captures Homologue-Resolved HP1 Phase Separation. J. Chem. Theory Comput. 2021, 17, 3134–3144.

(53) Latham, A. P. A.; Zhang, B. Improving Coarse-Grained Protein Force Fields with Small-Angle X-ray Scattering Data. J. Phys. Chem. B 2019, 123, 1026–1034.

(54) Latham, A. P.; Zhang, B. Maximum Entropy Optimized Force Field for Intrinsically Disordered Proteins. J. Chem. Theory Comput. 2019, 16, 773–781.

(55) Latham, A. P.; Zhang, B. Unifying coarse-grained force fields for folded and disordered proteins. Curr. Opin. Struct. Biol. 2022, 72, 63–70.

(56) Marrink, S. J.; Risselada, H. J.; Yefimov, S.; Tieleman, D. P.; De Vries, A. H. The MARTINI force field: Coarse grained model for biomolecular simulations. J. Phys. Chem. B 2007, 111, 7812–7824.

(57) Souza, P. C. et al. Martini 3: a general purpose force field for coarse-grained molecular dynamics. Nat. Methods 2021, 18, 382–388.

(58) Thomasen, F. E.; Pesce, F.; Roesgaard, M. A.; Tesei, G.; Lindorff-Larsen, K. Improving Martini 3 for Disordered and Multidomain Proteins. J. Chem. Theory Comput. 2022, 18, 2033–2041.

(59) Benayad, Z.; Von Bülow, S.; Stelzl, L. S.; Hummer, G. Simulation of FUS Protein Condensates with an Adapted Coarse-Grained Model. J. Chem. Theory Comput. 2021, 17, 525–537.

(60) Tsanai, M.; Frederix, P. W. J. M.; Schroer, C. F. E.; Souza, P. C. T.; Marrink, S. J. Coacervate formation studied by explicit solvent coarse-grain molecular dynamics with the Martini. Chem. Sci. 2021,

(61) Zhang, Y.; Feller, S. E.; Brooks, B. R.; Pastor, R. W. Computer simulation of liquid/liquid interfaces. I. Theory and application to octane/water. J. Chem. Phys. 1995, 103, 10252–10266.

(62) McDaniel, J. R.; Radford, D. C.; Chilkoti, A. A Unified Model for de Novo Design of Elastin-like Polypeptides with Tunable Inverse Transition Temperatures. Biomacromolecules 2013, 14, 2866–2872.

(63) Meyer, D. E.; Chilkoti, A. Quantification of the Effects of Chain Length and Concentration on the Thermal Behavior of Elastin-like Polypeptides. Biomacromolecules 2004, 5, 846–851.

(64) Roe, R. J. Theory of the interface between polymers or polymer solutions. I. Two components system. J. Chem. Phys. 1975, 62, 490–499.

(65) Ye, S.; Latham, A. P.; Tang, Y.; Hsiung, C.-H.; Chen, J.; Luo, F.; Liu, Y.; Zhang, B.; Zhang, X. Micropolarity governs the structural organization of biomolecular condensates. Nat. Chem. Biol. 2023, 1–9.

(66) Hassouneh, W.; Zhulina, E. B.; Chilkoti, A.; Rubinstein, M. Elastin-like Polypeptide Diblock Copolymers Self-Assemble into Weak Micelles. Macromolecules 2015, 48, 4183–4195.

(67) Bates, F. S.; Fredrickson, G. H. Block copolymers-designer soft materials. Phys. Today 1999, 52, 32–38.

(68) Mai, Y.; Eisenberg, A. Self-assembly of block copolymers. Chem. Soc. Rev. 2012, 41, 5969–5985.

(69) Zhulina, E. B.; Adam, M.; Larue, I.; Sheiko, S. S.; Rubinstein, M. Diblock copolymer micelles in a dilute solution. Macromolecules 2005, 38, 5330–5351.

(70) Statt, A.; Casademunt, H.; Brangwynne, C. P.; Panagiotopoulos, A. Z. Model for disordered proteins with strongly sequence-dependent liquid phase behavior. J. Chem. Phys. 2020, 152.

(71) Wesśen, J.; Das, S.; Pal, T.; Chan, H. S. Analytical Formulation and Field-Theoretic Simulation of Sequence-Specific Phase Separation of Protein-Like Heteropolymers with Short- and Long-Spatial-Range Interactions. J. Phys. Chem. B 2022, 126, 9222–9245.

(72) Liu, Y.; Fares, M.; Dunham, N. P.; Gao, Z.; Miao, K.; Jiang, X.; Bollinger, S. S.; Boal, A. K.; Zhang, X. AgHalo: A Facile Fluorogenic Sensor to Detect Drug-Induced Proteome Stress. Angew. Chemie - Int. Ed. 2017, 56, 8672–8676.

(73) Shen, B.; Jung, K. H.; Ye, S.; Hoelzel, C. A.; Wolstenholme, C. H.; Huang, H.; Liu, Y.; Zhang, X. A dual-functional BODIPY-based molecular rotor probe reveals different viscosity of protein aggregates in live cells. Aggregate 2022, 2–8.

(74) Chambers, J. E.; Kubánková, M.; Huber, R. G.; Ĺopez-Duarte, I.; Avezov, E.; Bond, P. J.; Marciniak, S. J.; Kuimova, M. K. An Optical Technique for Mapping Microviscosity Dynamics in Cellular Organelles. ACS Nano 2018, 12, 4398–4407.

(75) Regy, R. M.; Thompson, J.; Kim, Y. C.; Mittal, J. Improved coarse-grained model for studying sequence dependent phase separation of disordered proteins. Protein Sci. 2021,

(76) Wimley, W. C.; White, S. H. Experimentally determined hydrophobicity scale for proteins at membrane interfaces. Nat. Struct. Biol. 1996, 3, 842–848.

(77) Wimley, W. C.; Creamer, T. P.; White, S. H. Solvation energies of amino acid side chains and backbone in a family of host - Guest pentapeptides. Biochemistry 1996, 35, 5109–5124.

(78) Wilce, M. C.; Aguilar, M. I.; Hearn, M. T. Physicochemical Basis of Amino Acid Hydrophobicity Scales: Evaluation of Four New Scales of Amino Acid Hydrophobicity Coefficients Derived from RP-HPLC of Peptides. Anal. Chem. 1995, 67, 1210–1219.

(79) Zimmerman, J. M.; Eliezer, N.; Simha, R. The characterization of amino acid sequences in proteins by statistical methods. J. Theor. Biol. 1968, 21, 170–201.

(80) Radzicka, A.; Wolfenden, R. Comparing the Polarities of the Amino Acids: Side-Chain Distribution Coefficients between the Vapor Phase, Cyclohexane, 1-Octanol, and Neutral Aqueous Solution. Biochemistry 1988, 1664–1670.

(81) Lawson, E. Q.; Sadler, A. J.; Harmatz, D.; Brandau, D. T.; Micanovic, R.; MacElroy, R. D.; Middaugh, C. R. A simple experimental model for hydrophobic interactions in proteins. J. Biol. Chem. 1984, 259, 2910–2912.

(82) Simm, S.; Einloft, J.; Mirus, O.; Schleiff, E. 50 years of amino acid hydrophobicity scales: Revisiting the capacity for peptide classification. Biol. Res. 2016, 49, 1–19.

(83) Kapcha, L. H.; Rossky, P. J. A simple atomic-level hydrophobicity scale reveals protein interfacial structure. J. Mol. Biol. 2014, 426, 484–498.

(84) Tesei, G.; Schulze, T. K.; Crehuet, R.; Lindorff-larsen, K. Accurate model of liquid – liquid phase behavior of intrinsically disordered proteins from optimization of single-chain properties. Proc. Natl. Acad. Sci. U.S.A. 2021, e2111696118.

(85) Dannenhoffer-Lafage, T.; Best, R. B. A Data-driven Hydrophobicity Scale for Predicting Liquid-Liquid Phase Separation of Proteins. J. Phys. Chem. B 2021, 125, 4046–4056.

(86) Rose, G. D.; Geselowitz, A. R.; Lesser, G. J.; Lee, R. H.; Zehfus, M. H. Hydrophobicity of Amino Acid Residues in Globular Proteins. Science 1985, 229, 834.

(87) Tien, M. Z.; Meyer, A. G.; Sydykova, D. K.; Spielman, S. J.; Wilke, C. O. Maximum allowed solvent accessibilites of residues in proteins. PLoS ONE 2013, 8.

(88) Kyte, J.; Doolittle, R. F. A simple method for displaying the hydropathic character of a protein. J. Mol. Biol. 1982, 157, 105–132.

(89) Wei, M. T.; Elbaum-Garfinkle, S.; Holehouse, A. S.; Chen, C. C. H.; Feric, M.; Arnold, C. B.; Priestley, R. D.; Pappu, R. V.; Brangwynne, C. P. Phase behaviour of disordered proteins underlying low density and high permeability of liquid organelles. Nat. Chem. 2017, 9.

(90) Sticke, D. F.; Presta, L. G.; Dill, K. A.; Rose, G. D. Hydrogen bonding in globular proteins. J. Mol. Biol. 1992, 226, 1143–1159.

(91) Hofmann, H.; Soranno, A.; Borgia, A.; Gast, K.; Nettels, D.; Schuler, B. Polymer scaling laws of unfolded and intrinsically disordered proteins quantified with single-molecule spectroscopy. Proc. Natl. Acad. Sci. U.S.A. 2012, 109, 16155–16160.

(92) Dignon, G. L.; Zheng, W.; Best, R. B.; Kim, Y. C.; Mittal, J. Relation between single-molecule properties and phase behavior of intrinsically disordered proteins. Proc. Natl. Acad. Sci. U.S.A. 2018, 201804177.

(93) Wohl, S.; Jakubowski, M.; Zheng, W. Salt-Dependent Conformational Changes of Intrinsically Disordered Proteins. J. Phys. Chem. Lett. 2021, 12, 6684–6691.

(94) Welsh, T. J.; Krainer, G.; Espinosa, J. R.; Joseph, J. A.; Sridhar, A.; Jahnel, M.; Arter, W. E.; Saar, K. L.; Alberti, S.; Collepardo-Guevara, R.; Knowles, T. P. Surface Electrostatics Govern the Emulsion Stability of Biomolecular Condensates. Nano Lett. 2022, 22, 612–621.

(95) Martin, E. W.; Holehouse, A. S.; Peran, I.; Farag, M.; Incicco, J. J.; Bremer, A.; Grace, C. R.; Soranno, A.; Pappu, R. V.; Mittag, T. Valence and patterning of aromatic residues determine the phase behavior of prion-like domains. Science 2020, 367, 694–699.

(96) Zheng, W.; Dignon, G.; Brown, M.; Kim, Y. C.; Mittal, J. Hydropathy Patterning Complements Charge Patterning to Describe Conformational Preferences of Disordered Proteins. J. Phys. Chem. Lett. 2020, 11, 3408–3415.

(97) Huang, J.; Rauscher, S.; Nawrocki, G.; Ran, T.; Feig, M.; de Groot, B. L.; Grubmüller, H.; MacKerell, A. D. CHARMM36m: an improved force field for folded and intrinsically disordered proteins. Nat. Methods 2016, 14, 71–73.

(98) Berendsen, H. J.; van der Spoel, D.; van Drunen, R. GROMACS: A message-passing parallel molecular dynamics implementation. Comput. Phys. Commun. 1995, 91, 43– 56.

(99) Baek, M. et al. Accurate prediction of protein structures and interactions using a three-track neural network. Science 2021, 373, 871–876.

(100) Bonneau, R.; Tsai, J.; Ruczinski, I.; Chivian, D.; Rohl, C.; Strauss, C. E.; Baker, D. Rosetta in CASP4: Progress in ab initio protein structure prediction. Proteins 2001, 45, 119–126.

(101) Dignon, G. L.; Zheng, W.; Kim, Y. C.; Best, R. B.; Mittal, J. Sequence determinants of protein phase behavior from a coarse-grained model. PLoS Comput. Biol. 2018, 14, 1–23.

(102) Ladd, A. J.; Woodcock, L. V. Triple-point coexistence properties of the lennard-jones system. Chem. Phys. Lett. 1977, 51, 155–159.

(103) Tribello, G. A.; Giberti, F.; Sosso, G. C.; Salvalaglio, M.; Parrinello, M. Analyzing and Driving Cluster Formation in Atomistic Simulations. J. Chem. Theory and Comput. 2017, 13, 1317–1327.

(104) Gowers, R.; Linke, M.; Barnoud, J.; Reddy, T.; Melo, M.; Seyler, S.; Domański, J.; Dotson, D.; Buchoux, S.; Kenney, I.; Beckstein, O. MDAnalysis: A Python Package for the Rapid Analysis of Molecular Dynamics Simulations. Proceedings of the 15th Python in Science Conference. 2016.

(105) Michaud-Agrawal, N.; Denning, E. J.; Woolf, T. B.; Beckstein, O. MDAnalysis: a toolkit for the analysis of molecular dynamics simulations. J. Comput. Chem. 2011, 32, 2319–2327.

(106) Eisenhaber, F.; Lijnzaad, P.; Argos, P.; Sander, C.; Scharf, M. The Double Cubic Lattice Method: Efficient Approaches to Numerical Integration of Surface Area and Volume and to Dot Surface Contouring of Molecular Assemblies. J. Comput. Chem. 1995, 16, 273–284.

(107) Joosten, R. P.; Te Beek, T. A.; Krieger, E.; Hekkelman, M. L.; Hooft, R. W.; Schneider, R.; Sander, C.; Vriend, G. A series of PDB related databases for everyday needs. Nucleic Acids Res. 2015, 43, D364–D368.

(108) Kabsch, W.; Sander, C. Dictionary of Protein Secondary Structure: Pattern Recognition of Hydrogen-Bonded and Geometrical. Biopolymers 1983, 22, 2577–2637.

(109) Smith, P.; Ziolek, R. M.; Gazzarrini, E.; Owen, D. M.; Lorenz, C. D. On the interaction of hyaluronic acid with synovial fluid lipid membranes. Phys. Chem. Chem. Phys. 2019, 21, 9845–9857.

