## Supporting Information for "Microphase Separation Produces Interfacial Environment within Diblock Biomolecular Condensates"

#### Contents

**Supplemental Methods** **S4**

Condensate simulation details . . . . . . . . . . . . . . . . . . . . . . . . . . . . . S4

Homopolymer simulations . . . . . . . . . . . . . . . . . . . . . . . . . . . . S4

MOFF simulations . . . . . . . . . . . . . . . . . . . . . . . . . . . . . . . . S5

MARTINI simulations . . . . . . . . . . . . . . . . . . . . . . . . . . . . . . S7

Atomistic simulations . . . . . . . . . . . . . . . . . . . . . . . . . . . . . . S8

Monomer simulation details . . . S10
Experimental characterization of ELP condensates . . . S10
Plasmid synthesis . . . S11
Protein expression and purification . . . S11
Sequence dependence of micro-viscosity and polarity . . . S12

**Supplemental Theory** **S14**
Relationship between the transition temperature $T_t$ and the critical temperature $T_C$ S14
Relationship between the critical temperature $T_c$, the Flory-Huggins parameter $\chi$, and the surface tension, $\tau$ . . . S14
Relationship between the critical temperature, $T_c$, and hydrophobicity scales . . . S16

**Supplemental figures** **S17**
Figure S1 . . . S17
Figure S2 . . . S18
Figure S3 . . . S19
Figure S4 . . . S20
Figure S5 . . . S21
Figure S6 . . . S22
Figure S7 . . . S23
Figure S8 . . . S24
Figure S9 . . . S25
Figure S10 . . . S26
Figure S11 . . . S29
Figure S12 . . . S29
Figure S13 . . . S29
Figure S14 . . . S30
Figure S15 . . . S31

Figure S16 . . . . . . . . . . . . . . . . . . . . . . . . . . . . . . . . . . . . . S32

Figure S17 . . . . . . . . . . . . . . . . . . . . . . . . . . . . . . . . . . . . . S33

Figure S18 . . . . . . . . . . . . . . . . . . . . . . . . . . . . . . . . . . . . . S34

Table S1 . . . . . . . . . . . . . . . . . . . . . . . . . . . . . . . . . . . . . . S35

Table S2 . . . . . . . . . . . . . . . . . . . . . . . . . . . . . . . . . . . . . . S36

**Experimental sequences** **S37**

### Supplemental methods

#### Condensate simulation details

We employed a multiscale approach to study ELP condensates. Our simulations consisted of four steps with increasing resolution. We start with homopolymer and MOFF[S1] simulations that help generate initial configurations for the peptides within the condensate. Then we use the results from the explicit solvent coarse-grained model, MARTINI,[S2] and all atom explicit solvent force field, CHARMM36m,[S3] for further examination and quantitative analysis. Here, we describe the force field and simulation parameters at each step. Simulations were conducted using GROMACS.[S4] Homopolymer and MOFF configurations were performed on GROMACS 2018.4, which allowed for tabulated interactions not allowed in current versions of GROMACS. MARTINI and all atom simulations were performed using GROMACS 2021.

##### Homopolymer simulations

We used an identical homopolymer simulation to help generate starting configurations for elastin-like polypeptides. By using an identical configuration, all future simulations begin with the same molar density of protein, despite different sequences of ELPs. Our homopolymer force field was based on the MOFF force field. It assumed random coil secondary structure and non-bonded interactions identical to MOFF's valine-valine interaction. More specifically,

$$U_{\text{hom}}(\boldsymbol{r}) = U_{\text{bond}} + U_{\text{angle}} + U_{\text{contact}}, \tag{S1}$$

where $U_{\text{bond}}$, $U_{\text{angle}}$, and $U_{\text{contact}}$ account for the bonds between neighboring amino acids, angles between three neighboring amino acids, and nonbonded interactions between amino acids that are separated by more than 3 bonds, respectively.

The bonding potential $U_{\text{bond}} = \sum_i V_b(r_{i,i+1})$, where $i$ is the bead and

$$V_b(r_{i,i+1}) = \frac{k_b}{2}(r_{i,i+1} - r_0)^2. \tag{S2}$$

We used $k_b = 1000$ kJ mol$^{-1}$nm$^{-2}$ and $r_0 = 0.38$ nm. The angular potential $U_{\text{angle}} = \sum_i V_a(\theta_i)$ with

$$V_a(\theta_i) = \frac{k_a}{2}(\theta_i - \theta_0)^2. \tag{S3}$$

$\theta_i$ is the angle formed between the three consecutive beads $i$, $i+1$, and $i+2$. $k_a = 120$ kJ mol$^{-1}$ deg$^{-2}$, and $\theta_0 = 127$ degrees. The contact potential, $U_{\text{contact}} = \sum_{ij} V_{\text{nb}}(r_{ij}, \epsilon_{IJ})$. The pairwise potential is a combination of excluded volume and contact terms given by

$$V_{\text{nb}}(r_{ij}) = \frac{|\epsilon|\sigma^{12}}{r_{ij}^{12}} + \epsilon\mathcal{C}(r_{ij}). \tag{S4}$$

$i$ and $j$ correspond to interacting beads, $\epsilon = -2.327339$ kJmol$^{-1}$, and $\sigma = 0.586$ nm.

In this force field, we placed 40 polymers in a 75 nm by 75 nm by 75 nm simulation box. After steepest decent energy minimization, we performed an NPT simulation for 0.1 $\mu$s at 150 K and 1 bar, using a Parrinello-Rahman isotropic bariostat and time coupling constant of 1 ps. This resulted in a configuration of forty homopolymers compressed into a 6.3782 nm by 6.3782 nm by 6.3782 nm simulation box.

#### MOFF simulations

From the starting configuration produced by the homopolymer simulation, we wanted to relax the contacts to be representative of those with ELPs before we began our explicit solvent simulations. For this equilibration step, we turned to the MOFF force field. ELPs are known to have minimal secondary structure in both the dense and dilute state,[S5,S6] so we assumed the secondary structure to be similar to that of a random coil. Under these simplifications, MOFF can be specified as

$$U_{\text{hom}}(\boldsymbol{r}) = U_{\text{bond}} + U_{\text{angle}} + U_{\text{electrostatics}} + U_{\text{contact}}, \tag{S5}$$

where $U_{\text{bond}}$ is described by Eq. S2, $U_{\text{angle}}$ is described by Eq. S3, $U_{\text{electrostatics}}$ accounts for electrostatic interactions between charged amino acids, and $U_{\text{contact}}$ accounts for non-bonded interactions.

$U_{\text{electrostatics}}$ accounts for electrostatic interactions between charged residues. Based on the Debye-Hückel theory, it can be approximated as

$$U_{\text{electrostatics}} = \sum_{i,j} \frac{q_i q_j}{4\pi\epsilon_0 r_{ij}\epsilon(r_{ij})} \exp(-r_{ij}/\lambda_D), \tag{S6}$$

where $\epsilon_0$ is the permittivity of free space, $\lambda_D$ is the Debye screening length, $r_{ij}$ is the distance between particles $i$ and $j$, and $q_i$ and $q_j$ are charges of particles $i$ and $j$. Charges of individual residues can be found in Table S1 of of Latham and Zhang.[S1] We used a distance dependent dielectric constant, $\epsilon(r_{ij})$, to capture the change in the solvation environment upon protein folding.[S7,S8] This dielectric takes the form

$$\epsilon(r_{ij}) = A + \frac{B}{1 + ke^{-\lambda B r_{ij}}}, \tag{S7}$$

where $A = -8.5525$, $k = 7.7839$, $\lambda = 0.03627\text{nm}^{-1}$, and $B = \epsilon_{\text{w}} - A$, where $\epsilon_{\text{w}} = 78.4$ is the dielectric constant of water.

The contact potential, $U_{\text{contact}} = \sum_{ij} V_{\text{nb}}(r_{ij}, \epsilon_{IJ})$. The pairwise potential is a combination of excluded volume and contact terms given by

$$V_{\text{nb}}(r_{ij}, \epsilon_{IJ}) = \frac{|\epsilon_{IJ}|\sigma_{IJ}^{12}}{r_{ij}^{12}} + \epsilon_{IJ}\mathcal{C}(r_{ij}). \tag{S8}$$

$I$ and $J$ correspond to the amino acid type of bead $i$ and $j$. $\sigma_{IJ} = \frac{\sigma_I + \sigma_J}{2}$ is defined using the individual size of each amino acid type, and full parameters for the implementation of this potential can be found in Table S7 and Table S8 of Latham and Zhang.[S1]

In this force field, we conducted energy minimization, and then a 0.1 $\mu$s NVT simulation, with a timestep of 10 fs. During this simulation, we utilize the GROMACS simulated annealing protocol to linearly raise the temperature from 150 K to 300 K, with a time coupling constant of 100 ps.

#### MARTINI simulations

The final implicit solvent, coarse grained MOFF configuration was used as a starting point for our explicit solvent coarse-grained simulations with the MARTINI force field. To convert our $\alpha-$carbon only configuration to a MARTINI representation, we converted the $\alpha-$carbon snapshot to an all atom representation using tleap, which is part of AMBER tools 2020.[S9] Next, we converted the all atom configuration to a MARTINI3 configuration using martinize2.[S2] We used random coil secondary structure throughout the entire peptide and enforced neutral termini for each peptide chain. We then expanded the Z-dimension of the simulation box to 40 nm and centered the protein in the simulation box. All simulations thus started with box dimensions of 6.3782 nm by 6.3782 nm by 40.0000 nm. This change in box dimensions was preparation for slab simulations, which enabled us to probe the physical properties of the condensate.[S10] In this setup, the high-density protein phase became a dense slab that was periodic in $x$ and $y$, but did not interact with its periodic image in $z$ due to the size of the box. The lack of periodic interactions in the $z$ direction allowed the formation of two interfaces between a dense biopolymer phase and the solvent phase. We then solvated the peptide with MARTINI water and 1 M NaCl.

We started with steepest decent energy minimization. Then, we performed NVT equilibration with a timestep of 10 fs for a total simulation time of 5 ns. The protein and solvent were held at 313.15 K by separately coupling them to a velocity-rescaling thermostat with a time coupling constant of 1 ps. We then equilibrated our simulations in an NPT ensemble for 10 ns with a timestep of 20 fs. We separately coupled protein and solvent to a temperature of 313.15 K with a velocity-rescaling thermostat with a time coupling constant of 1 ps. The pressure was fixed using a Berendsen bariostat with a reference pressure of 1 bar, compressibility of $4.5 \times 10^{-5}$ bar$^{-1}$, and a time coupling constant of 5 ps. We then performed our production simulation in an NPT ensemble with semiisotropic pressure coupling, which is sometimes called an $NP_NAT$ ensemble.[S11] This technique fixes the pressure normal to the protein surface, which allows us to calculate the surface tension of the protein-solvent interface. Temperature was coupled separately to 313.15 K for the protein and solvent using a velocity-rescaling thermostat with a time coupling constant of 1 ps. A Parrinello-Rahman semiisotropic bariostat was used to couple pressure at a time coupling constant of 10 ps. A compressibility of $4.5 \times 10^{-5}$ $\text{bar}^{-1}$ and pressure of 1 bar was used in the Z-dimension, but the compressibility was set to 0 in the X-Y dimensions. The production simulations lasted for 100 $\mu$s and used a timestep of 20 fs. The first 25 $\mu$s was discarded for equilibration, and configurations were recorded every 1000 ps for analysis.

Finally, not every $\alpha-$carbon, implicit solvent representation from MOFF could be stably converted into a higher resolution, explicit solvent MARTINI representation. If energy minimization was unable to resolve clashes within the MARTINI configuration we converted from the final step in the MOFF simulation, we iteratively moved back one snapshot and repeated the procedure until a stable configuration was found.

We note that many studies of IDPs with MARTINI have scaled the protein-water interactions to improve agreement between simulation and experiment.[S12,S13] However, in the case of ELPs studied here, we find the surface tensions given by the default MARTINI model are reasonable based on experimentally informed theories of ELP condensates[S14] and experimental observations of surface tension in other biological condensates,[S15] which suggest biological condensates may have interfacial tensions as large as ~1 mN/m.

##### Atomistic simulations

Proteins were described using the CHARMM36m force field.[S3] Proteins were capped using acetylated N-terminus and amidated C-terminus. TIP3P water was used, and the water-hydrogen Lennard-Jones well depth was scaled to -0.10 kcal/mol. This scaling has been shown to better describe IDPs in previous work.[S3]

All atom simulations were started based on the final timestep of MARTINI simulations. MARTINI configurations were converted to CHARMM36m configurations using the backward script.[S3,S16] This code converts MARTINI configurations into all-atom representations and then ensures configuration stability through a series of short energy minimization and restrained molecular dynamics simulations. The final output for these configurations was utilized as the starting point for all atom simulations. If the backward script did not result in as stable all-atom representation, we iteratively moved back one snapshot in the MARTINI and repeated the backward procedure until a stable all atom initial configuration was found. We performed an additional round of energy minimization, and then equilibrated the protein in the NVT ensemble for 100 ps with a timestep of 1 fs. We used a Nose-Hoover thermostat to separately couple the protein and solvent to 313.15 K with a time coupling constant of 1.0 ps. Position restraints were applied with a force constant of 400 $\text{kJmol}^{-1}\text{nm}^{-2}$ for the backbone and 40 $\text{kJmol}^{-1}\text{nm}^{-2}$ for the side chains. We then performed NPT equilibration for 5 ns with a timestep of 2 fs. We used a Nose-Hoover thermostat to separately couple the protein and solvent to 313.15 K with a time coupling constant of 1.0 ps, and coupled the pressure using an isotropic Parrinello-Rahman bariostat with a compressibility of $4.5 \times 10^{-5}$ $\text{bar}^{-1}$, a reference pressure of 1 bar, and a time coupling constant of 2 ps. We then performed an additional equilibration in the $\text{NP}_\text{N}\text{AT}$ ensemble by using a semiisotropic bariostat for 5 ns with a timestep of 2 fs. We used a Nose-Hoover thermostat to separately couple the protein and solvent to 313.15 K with a time coupling constant of 1.0 ps. We coupled the pressure using an semiisotropic Parrinello-Rahman bariostat with a compressibility of $4.5 \times 10^{-5}$ $\text{bar}^{-1}$ and a reference pressure of 1 bar in the Z-dimension, but set the compressibility to 0 in the X-Y dimensions. The pressure-time coupling constant was 2 ps. We then performed production runs in the $\text{NP}_\text{N}\text{AT}$ ensemble. Like the previous simulation, we used a Nose-Hoover thermostat to separately couple the protein and solvent to 313.15 K with a time coupling constant of 1.0 ps, and coupled the pressure using an semiisotropic Parrinello-Rahman bariostat with a compressibility of $4.5 \times 10^{-5}$ $\text{bar}^{-1}$ and a reference pressure of 1 bar in the Z-dimension, but set the compressibility to 0 in the X-Y dimensions. The pressure-time coupling constant was 2 ps. Simulations lasted for 250 ns, and the first 50 ns was discarded for equilibration. We recorded timesteps every 10 ps for analysis.

#### Monomer simulation details

In addition to the condensates, we carried out atomistic simulations of ELP monomers for comparison. These simulations were conducted on all 20 sequences and began from RosettaFold predictions of the peptide structure.[S17,S18] RoseTTAFold was used in all cases except for $V_5M_5$, where not enough homologs were available for a prediction.[S17] In this case, Rosetta ab initio predictions were used instead.[S18] The force field was identical to the condensate simulations, and thus used the CHARMM36m force field, with shifted water potentials and capped termini.[S3] Each peptide was placed in a simulation box of 8 nm by 8 nm by 8 nm, which gives a $> 99\%$ chance of the side of the simulation box being longer than the end-to-end distance of an ideal chain with a Kuhn length of 0.55 nm, as has has been suggested for IDPs.[S19] Peptides were solvated in water with an ionic strength of 1 M.

After solvating each protein, we conducted energy minimization. Then, we performed 100 ps of equilibration with a timestep of 1 fs in the NVT ensemble. We used a Nose-Hoover thermostat to separately couple the protein and solvent to 313.15 K with a time coupling constant of 1.0 ps. Position restraints were applied with a force constant of 400 $\mathrm{kJmol^{-1}nm^{-2}}$ for the backbone and 40 $\mathrm{kJmol^{-1}nm^{-2}}$ for the side chains. Next, we performed NPT equilibration for 5 ns with a timestep of 2 fs. Simulations were conducted at 313.15 K and 1.0 bar. Finally, we conducted a production simulation in the NPT ensemble for 1 $\mu$s with a timestep of 2 fs, at 313.15 K and 1.0 bar. We excluded the first 100 ns of this production run for additional equilibration, which left 900 ns for analysis.

#### Experimental characterization of ELP condensates

##### Plasmid synthesis

Cloning vector pET-28a+ was modified to contain endonuclease recognition sites of BseRI, AcuI, and BglI for inserting the designed ELP sequence, similar to McDaniel et. al.[S20] Modified pET-28a+ was linearized using BseRI (NEB, China) and dephosphorylated with rSAP (NEB, China) for ELP assembly. ssDNA encoding for five pentapeptides (5-mer) sequences was ordered through GeneWiz, China. Then the ssDNA chains were annealed and phosphorylated to form double-stranded duplexes with sticky overhangs. Linearized pET-28a+ vectors and duplexes were ligated by the T4 DNA ligase (NEB,China). After obtaining 5-mer ELP plasmids, we performed the iterative cloning steps to increase the repeats through recursive directional ligation by the plasmid reconstruction method[S20] until achieving the target length of ELP. The sequences of all plasmids created at each step were confirmed by DNA sequencing.

##### Protein expression and purification

The targeted ELP plasmids were transformed into E. coli BL21 (DE3) cells. Cells were incubated overnight on the LB plate with Kanamycin. On the second day, a single colony was inoculated into 5 mL Luria Broth (LB) and serially diluted (1:100) into four different starter flasks (A-D). Starter cultures were shaken for 12-14 hours at 37°C, and the culture with the OD600 0.6 to 0.8 was inoculated into 1 L Terrific Broth (TB) containing 50 $\mu$g/mL kanamycin at 1:100 dilution. This culture was grown at 37°C and 220 rpm until the media OD600 reached 0.8, and the culture was then induced to express the protein with 0.5 mM isopropyl-$\beta$-D-thiogalactoside (IPTG). The cultures were then kept shaking for 20-24 hours at 37°C and 200 rpm before harvesting. The cells were collected by centrifuging at 6000 rpm at 4°C for 10 mins and resuspended in 1x PBS buffer (pH 7.4). The proteins were released from the cells by sonicating on ice for 6 min, with 2 s of pulsing followed by 8 s of resting on ice. The lysate was centrifuge at 14000 rpm at 4°C for 40 min. The supernatant was then collected and mixed with polyethyleneimine to a final concentration of 0.5 w/v% to precipitate nucleic acids. The mixture was then centrifuged at 14000 rpm and 4°C for 40 min. The supernatant was collected and added with NaCl(s) to get the final concentration of 4.5 M NaCl. The mixture was then shaken at 37°C to precipitate ELP and centrifuged at 10000 rpm at 37°C for 40 min. The pellet was saved and resuspended in double-distilled $H_2O$. This mixture was then shaken on ice to dissolve the ELP. The inverse transition cycling was repeated 1-2 more times.[S21] The protein solution was dialyzed in a 10 kDa membrane (SnakeSkin, Thermo Fischer Scientific) against 2 L double distilled $H_2O$ at 4°C, changing the dialysis water one time. The protein purity was determined by 12% SDS-PAGE gel electrophoresis and stained with 0.5 M copper chloride. ELP proteins were then concentrated and lyophilized to remove the water.

**Sequence dependence of micro-viscosity and polarity**

All ELP sequences were expressed in bacteria E. coli and purified to more than 95% purity, as analyzed by SDS-PAGE (Fig. S16 in the Supporting Information). The purified ELPs were subjected to mass spectroscopy measurement to confirm their identity (Table S2 and Fig. S17 in the Supporting Information). Using turbidity as a signal to report on the formation of liquid droplets as a function of temperature change, we found that $V_{30}A_{30}$ and $A_{30}V_{30}$ exhibited the same $T_t$ (transition temperature for phase separation) values (Fig. S8A-B in the Supporting Information), suggesting that both sequences underwent almost identical intermolecular interactions to transit from the one-liquid diffusive phase to the two-liquid phase separated phase. The same is true for $V_{30}G_{30}$ and $G_{30}V_{30}$ (Fig. S8C-D in the Supporting Information). To measure the viscosity and polarity of different micro-organizations of blocks, we used the established microenvironment-sensitive fluorogenic probes – boron dipyrromethene (BODIPY) and 7-sulfonamide benzoxadiazole (SBD). Based on previous reports, the fluorescence lifetime ($\tau$) of BODIPY would increase with elevating micro-viscosity;

whereas, the $\tau$ of SBD would rise with decreasing micro-polarity. The correlation of their $\tau$ values can be quantitatively determined using standard solvents, thus providing equations to quantify viscosity ($\eta$) and polarity (as $\epsilon$, dielectric constants) via the measurement of $\tau$ values (Fig. S18 in the Supporting Information).

Since we used $V_{30}X_{30}$ and $X_{30}V_{30}$ to quantify the V- and X-end of the V-X blocks, it is possible that the observed differences arose from the innate property of the $V_{30}X_{30}$ and $X_{30}V_{30}$ sequences. To rule out this artifact, we formed the ELP condensates with sequences of $V_{30}X_{30}$, $X_{30}V_{30}$, or the $V_{30}X_{30}$ and $X_{30}V_{30}$ mixture. The condensates were subsequently treated with the aldehyde-BODIPY and methyl-ester SBD fluorophores without the NHS ester reactive warhead (Fig. S9A in the Supporting Information). After brief incubation, aldehyde-BODIPY and methyl-ester SBD fluorophores were recruited into and homogeneously distributed in the ELP condensates. The fluorescence lifetime of aldehyde-BODIPY was the same for $V_{30}A_{30}$ (4.96 ns), $A_{30}V_{30}$ (4.99 ns), and their mixture (4.98 ns) (Fig. S9B in the Supporting Information, upper panel). Interestingly, this value is around the average (4.89 ns) of the A-end (4.35 ns) and the V-end (5.43 ns) labeled NHS-BODIPY. For the SBD measurement, methyl-ester SBD resulted in almost identical lifetime values of $V_{30}A_{30}$ (8.25 ns), $A_{30}V_{30}$ (8.27 ns), and their mixture (8.28 ns) (Fig. S9B in the Supporting Information, lower panel), again around the average values (7.88 ns) of the A-end (7.00 ns) and the V-end (8.75 ns) labeled NHS-SBD. In addition to the V-A blocks, similar observations were made for the V-G blocks as $V_{30}G_{30}$ and $G_{30}V_{30}$ sequences (Fig. S9C in the Supporting Information). The slight difference between the results is attributed to the experiment errors. Because the fluorophores did not covalently label the amino-terminus of the ELP peptides, their lifetime reports closer to the averaged property of the condensates instead of the microscopic property of the V-end or the X-end when the number of molecules is sufficient and the molecular distribution has no preference. Our results reveal that the $V_{30}X_{30}$ and $X_{30}V_{30}$ condensates exhibited similar macroscopic viscosity or polarity, suggesting that the previously observed different viscosity or polarity of $V_{30}X_{30}$ and $X_{30}V_{30}$ could be attributed to the microscopic property of the V-end or X-end.

### Supplemental Theory

#### Relationship between the transition temperature $T_t$ and the critical temperature and $T_C$

The temperature measured by Urry et al. and presented in the main text corresponds to the transition temperature, $T_t$.[S22] This temperature is defined as the value above which ELP forms an insoluble coacervate phase. The value of $T_t$ depends on the concentration of the polymer solution.

As shown by Chilkoti and coworkers,[S23,S24] the critical temperature $T_c$ is indeed linearly related to $T_t$ with the following relationship

$$T_t = T_c + \frac{k}{\text{length}} \ln \frac{C_c}{\text{conc}}. \tag{S9}$$

The above equation highlights the dependence of $T_t$ on the chain length (length) and polymer concentration (conc). The parameter $C_c$ is the corresponding theoretical polypeptide concentration that would be required to achieve $T_c$ and $k$ is the proportionality constant.

#### Relationship between the critical temperature $T_c$, the Flory-Huggins parameter $\chi$, and the surface tension, $\tau$

The Flory-Huggins parameter, $\chi$, is defined as

$$\chi = \frac{z}{2k_BT}[\epsilon_{\text{pp}} + \epsilon_{\text{ss}} - 2\epsilon_{\text{ps}}] = \frac{z\Delta\epsilon}{k_BT}, \tag{S10}$$

where $\Delta\epsilon = \frac{1}{2}[\epsilon_{\text{pp}} + \epsilon_{\text{ss}} - 2\epsilon_{\text{ps}}]$. $z$ is the coordination number, $T$ is the temperature, $k_B$ is the Boltzmann constant, and $\epsilon_{\text{pp}}$ are the strength of polymer-polymer, solvent-solvent, and polymer-solvent interactions respectively.[S25]

From the original derivation of Flory-Huggins theory, it can be shown that phase separation occurs when $\chi$ is greater than its critical value, or $\chi > \chi_c \approx \frac{1}{2}$. From $\chi_c$, we can derive the critical temperature as

$$T_c = \frac{2z\Delta\epsilon}{k_B}. \tag{S11}$$

Therefore, $\chi = \frac{T_c}{2T}$.

Relationships between the Flory Huggins parameter, $\chi$, and interfacial tension ($\tau$) have been investigated, and the relationship can be approximated as

$$\tau \propto \chi^{\alpha}, \tag{S12}$$

where $\alpha$ is a positive constant, whose exact value depends on the proximity of $\chi$ to the critical value of $\chi$ necessary for phase separation ($\chi_C$).[S26,S27] Correspondingly, we have $\tau \propto T_c^{\alpha}$.

We note that the expression for $\chi$ (Eq. S10) is simplified and assumes that $\Delta\epsilon$ is independent of temperature. However, changes in solvent packing upon phase separation can result in entropic contributions.[S28] In general, $\Delta\epsilon$ should be expressed as $\Delta\epsilon = \Delta h + T\Delta s$. The entropic contributions are essential for giving rise to phase separations with lower critical temperatures. Therefore, in the generalized case, we have

$$\chi = \frac{z\Delta\epsilon}{k_BT} = \frac{z(\Delta h + T\Delta s)}{k_BT}. \tag{S13}$$

At the critical temperature, we have

$$\frac{1}{2} = \frac{z(\Delta h + T_C\Delta s)}{k_BT_C}, \tag{S14}$$

and

$$T_C = \frac{z\Delta h}{k_B/2 - z\Delta s}. \tag{S15}$$

For systems exhibiting lower critical solution temperatures (LCST), $\Delta h < 0$ and $\Delta s > 0$. For $T_c > 0$, we have $k_B/2 < z\Delta s$.

Plugging the above expression back to Eq. S13, we have

$$\chi = \frac{k_B T_C/2 - z\Delta s T_C + zT\Delta s}{k_B T} = \frac{T_C}{T}\left(\frac{1}{2} - \frac{z\Delta s}{k_B}\right) + \frac{z\Delta s}{k_B}. \tag{S16}$$

Furthermore, as shown by Schauperl et al.[S29], $\Delta s$, while significant, remains relatively constant across different amino acids. Correspondingly, we have $\tau \propto \left[\frac{T_c}{T}\left(\frac{1}{2} - \frac{z\Delta s}{k_B}\right) + \frac{z\Delta s}{k_B}\right]^\alpha$ holds true in the more general case as well. Since $\frac{z\Delta s}{k_B} > \frac{1}{2}$, the slope between $\tau$ and $T_C$ is negative.

#### Relationship between the critical temperature, $T_c$ and hydrophobicity scales

As shown in Eq. S11, in the simple case $T_c$ is linearly related to $\Delta\epsilon$. For the more general case, from Eq. S15, we have

$$T_C = \frac{z\Delta h}{k_B/2 - z\Delta s} = \frac{z\Delta\epsilon}{k_B/2 - z\Delta s} - \frac{zT\Delta s}{k_B/2 - z\Delta s}. \tag{S17}$$

Assuming that $\Delta s$ is relatively independent of amino acids, as shown by Schauperl et al.[S29], $T_c$ is again linearly proportional to $\Delta\epsilon$.

$\Delta\epsilon = \frac{1}{2}[\epsilon_{\text{pp}} + \epsilon_{\text{ss}} - 2\epsilon_{\text{ps}}]$ can indeed be interpreted as the free energy cost of transferring a polymer bead from a solution phase to a polymer phase. It corresponds to the change of energy from a mixed state, with contacts between polymer and solvent ($\epsilon_{\text{ps}}$), to the demixed state with only polymer-polymer ($\epsilon_{\text{pp}}$) and solvent-solvent ($\epsilon_{\text{ss}}$) contacts.

Therefore, the transfer free energy, and the interactions among amino acids of ELPs, are expected to correlate with the critical temperature.

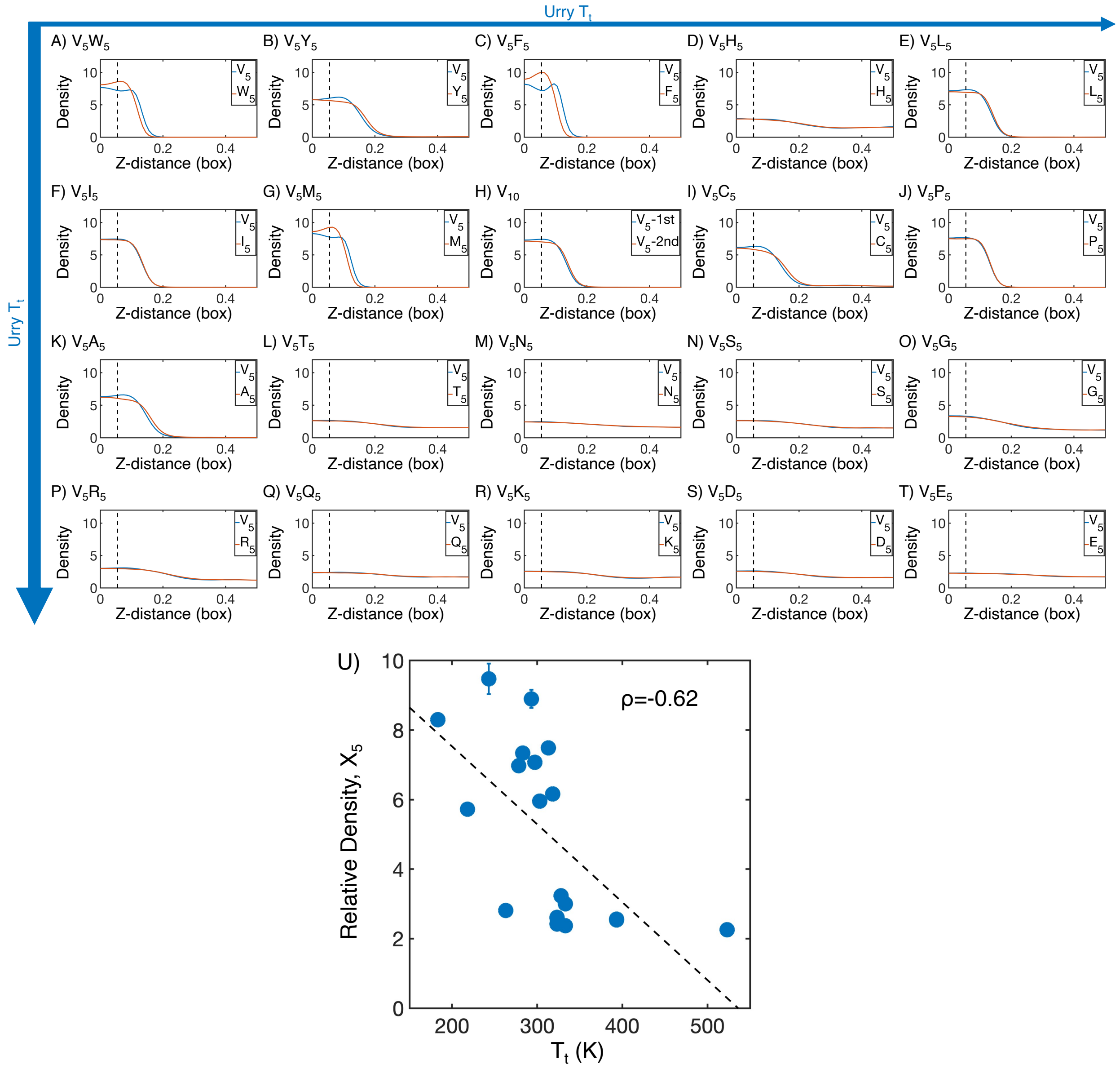


Figure S1: Mass density profiles of ELP condensates from MARTINI simulations. (A-T) The relative mass density along the $Z$-distance from the condensate center is shown for the V-substituted and X-substituted halves of each condensate. The $Z-$axis is defined as the direction perpendicular to the condensate-water interface. The dashed line represents a $Z$-distance of 0.06 box lengths away from the condensate center. Average density values below this threshold are used for correlation analysis in part U. Proteins are arranged by increasing $T_t$ (decreasing hydrophobicity) from Urry,[S22] with (A, $V_5W_5$) being the most hydrophobic and (T, $V_5E_5$) being the least hydrophobic. (U) Correlation between the mass fraction of the $X_5$ half of the condensate and transition temperature ($T_t$) from Urry.[S22] $\rho$ is the Pearson correlation coefficient between the two data sets, and the dashed diagonal line is the best fit line. Error bars represent standard deviations of the mean taken over 6 equally spaced box length intervals.

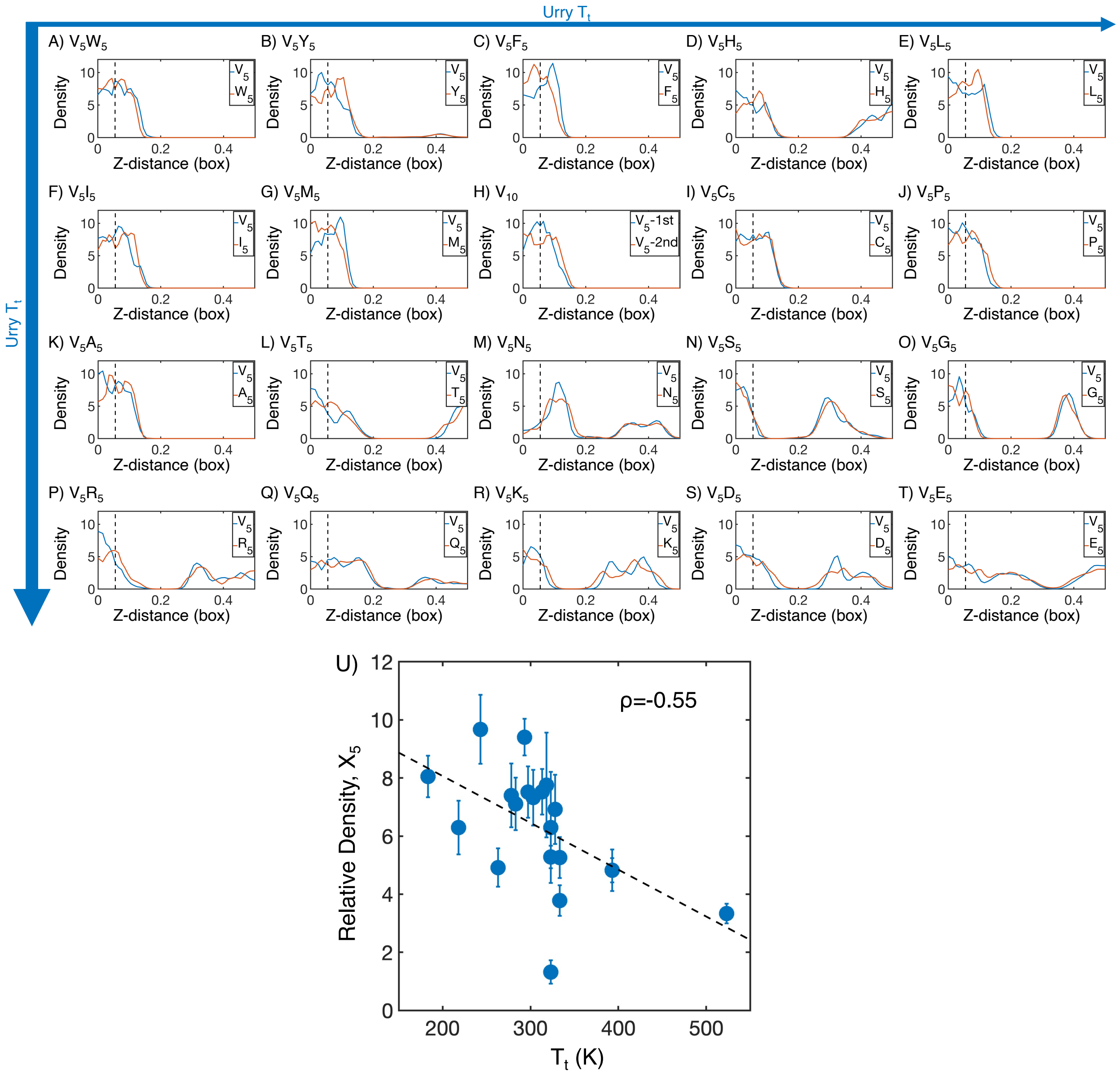

Figure S2: Mass density profiles of ELP condensates from all-atom simulations. (A-T) The relative mass density along the $Z$-distance from the condensate center is shown for the V-substituted and X-substituted halves of each condensate. The $Z-$axis is defined as the direction perpendicular to the condensate-water interface. The dashed line represents a $Z$-distance of 0.06 box lengths away from the condensate center. Average density values below this threshold are used for correlation analysis in part U. Proteins are arranged by increasing $T_t$ (decreasing hydrophobicity) from Urry,[S22] with (A, $V_5W_5$) being the most hydrophobic and (T, $V_5E_5$) being the least hydrophobic. (U) Correlation between the mass fraction of the $X_5$ half of the condensate and transition temperature ($T_t$) from Urry.[S22] $\rho$ is the Pearson correlation coefficient between the two data sets, and the dashed diagonal line is the best fit line. Error bars represent standard deviations of the mean taken over 6 equally spaced box length intervals.

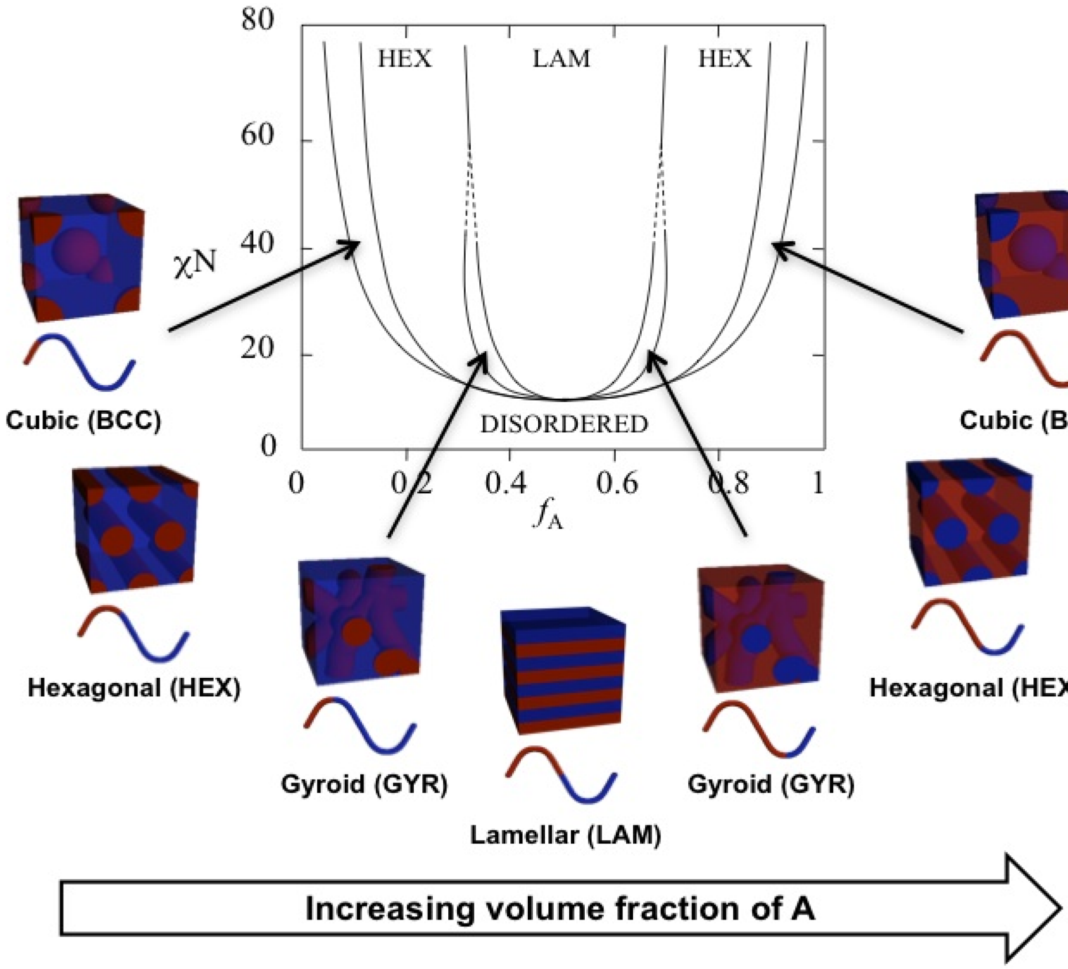


Figure S3: Theoretical phase diagram[S30] and corresponding morphologies for diblock copolymers. The phases are labeled as: body centered cubic (BCC), hexagonal cylinders (HEX), gyroid (GYR), and lamellar (LAM). $f_A$ is the volume fraction of a single polymer block, denoted A, $\chi$ is the Flory-Huggins interaction parameter, and $N$ is the total degree of polymerisation. Figure reproduced from ref.[S31] CC BY 4.0.

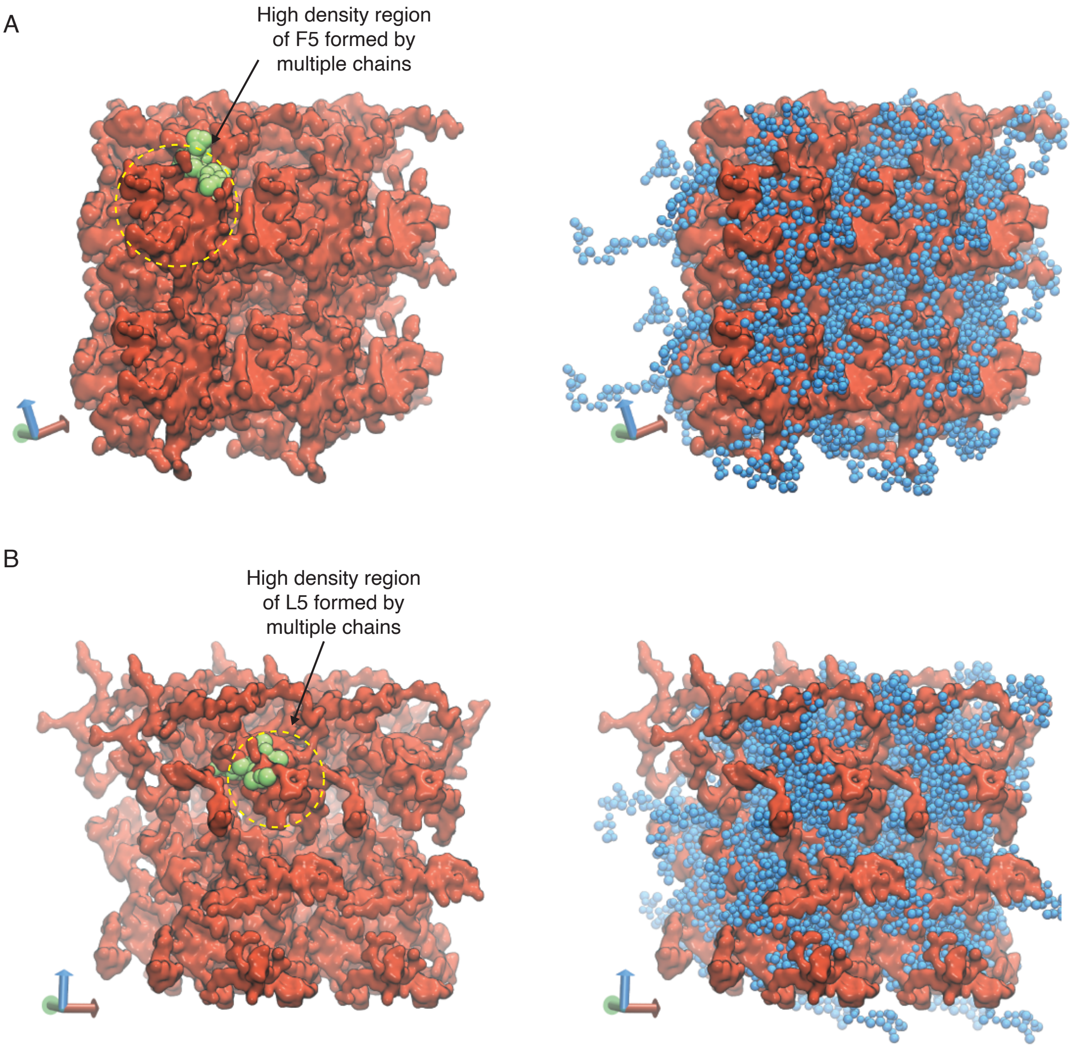


Figure S4: Representative configurations of (A) $V_5F_5$ and (B) $V_5L_5$ condensates from MARTINI simulations. The valine substituted half of the chain is colored blue ($V_5$) and the X substituted half of the chain is colored red ($X_5$). To highlight the interpenetrating networks formed by the two halves, only the X substituted half of the chain is shown on the left. Simulation interfaces are once repeated periodically in the positive x and positive y dimensions for clarity. High density regions formed by the multiple X substituted half of the chains are highlighted in yellow circles, with one of the chain shown in green.

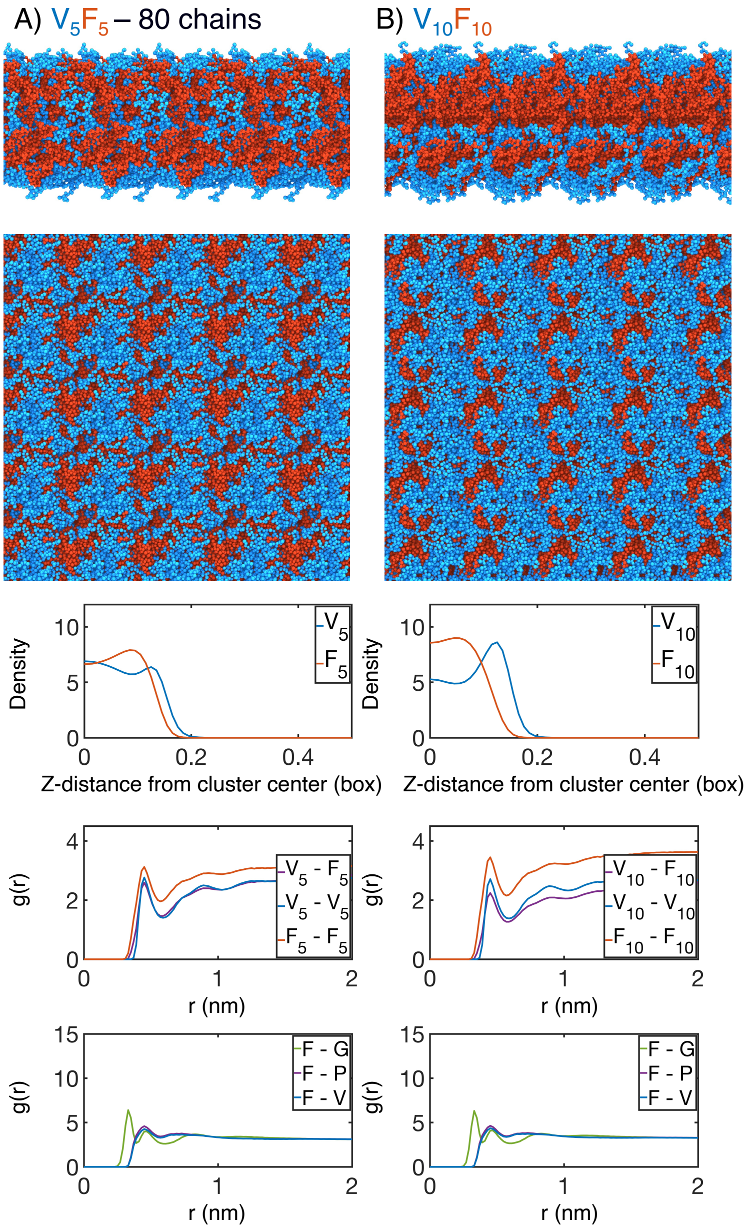


Figure S5: Finite size effect checks of our simulation methodology by (A) doubling the number of chains in the simulation box ($V_5F_5$ - 80 chains) or by (B) doubling the length of the peptide sequence ($V_{10}F_{10}$). In each case, the uppermost panel is a periodic image of the simulated condensate. Only the protein is shown, with red highlighting the X-substituted half of the condensate and blue highlighting the V-substituted half of the condensate. The second panel from the top is condensate viewed from the simulation interface. The middle panel is the mass density from the condensate center for for the V-substituted and X-substituted halves of each condensate. The second panel from the bottom is the radial distribution function g(r) for inter-chain coarse-grained beads, divided between the $V_5$ half of the chain and $X_5$ half of the chain. The bottom panel is the overall radial distribution function from the guest amino acid ($X_5$), to those amino acids native to the ELP sequence.

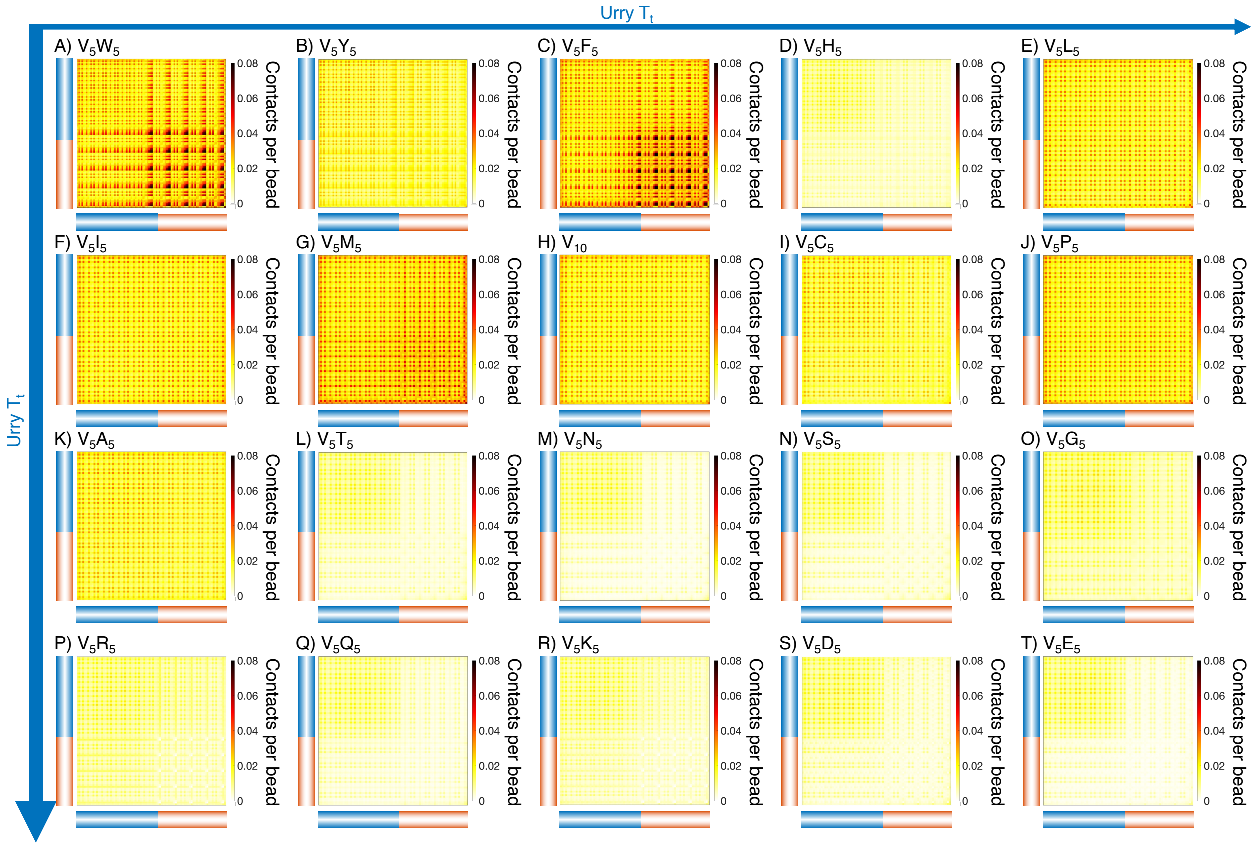


Figure S6: Intermolecular contact maps for ELP peptides from MARTINI simulations. The blue bar highlights the $V_5$ half of the condensate, while red bar highlights the $X_5$ half. Proteins are arranged by increasing $T_t$ (decreasing hydrophobicity) from Urry,[S22] with (A, $V_5W_5$) being the most hydrophobic and (T, $V_5E_5$) being the least hydrophobic.

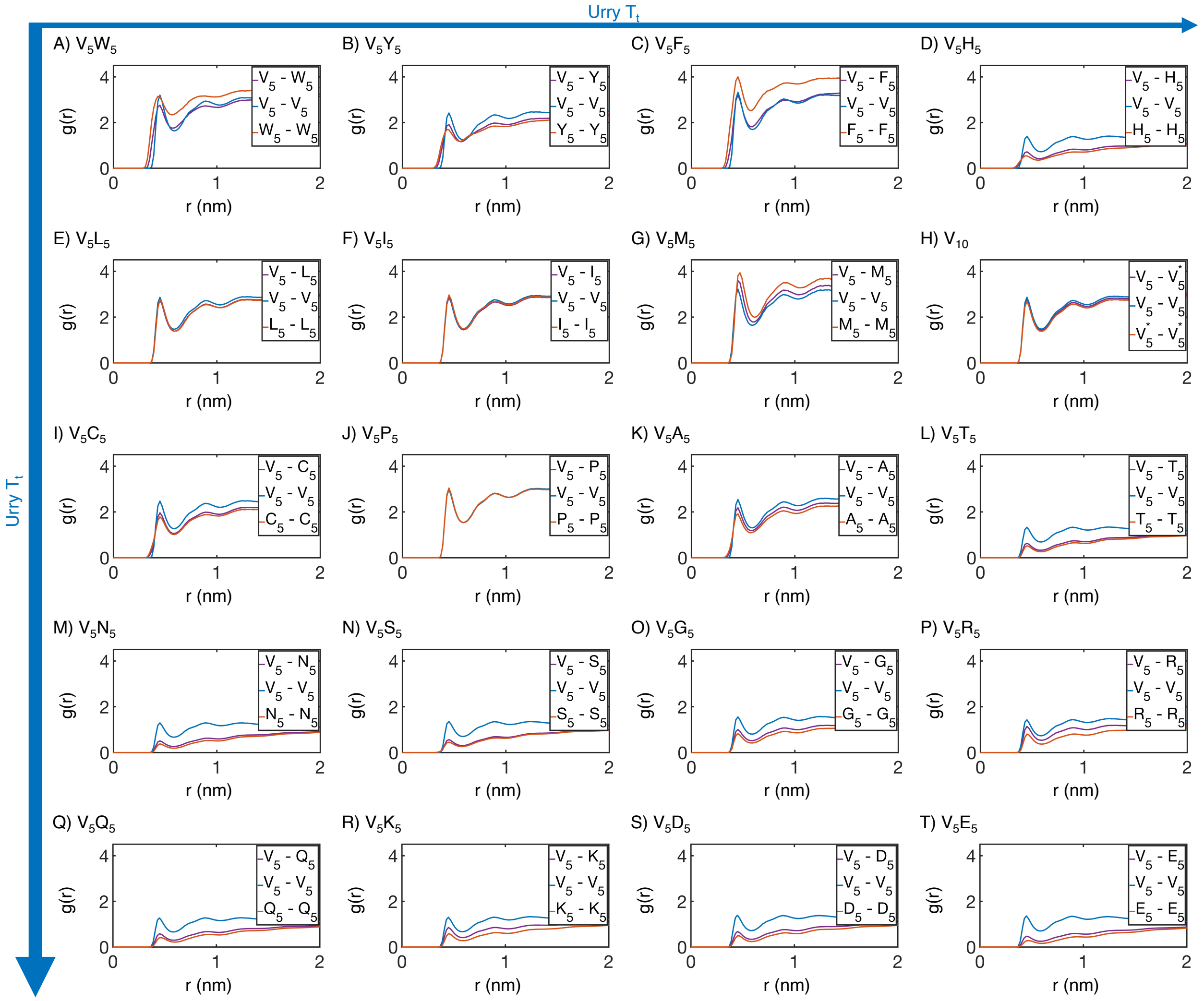


Figure S7: Radial distribution functions $g(r)$ from MARTINI simulations support the microphase separation of different ELP blocks. Amino acids from the $V_5$ and $X_5$ half of the ELP peptide are partitioned into two blocks, and we used distances within and across the blocks to compute $g(r)$. Distances between amino acids from the same peptide are excluded from computations. Proteins are arranged by increasing $T_t$ (decreasing hydrophobicity) from Urry,[S22] with (A, $V_5W_5$) being the most hydrophobic and (T, $V_5E_5$) being the least hydrophobic.

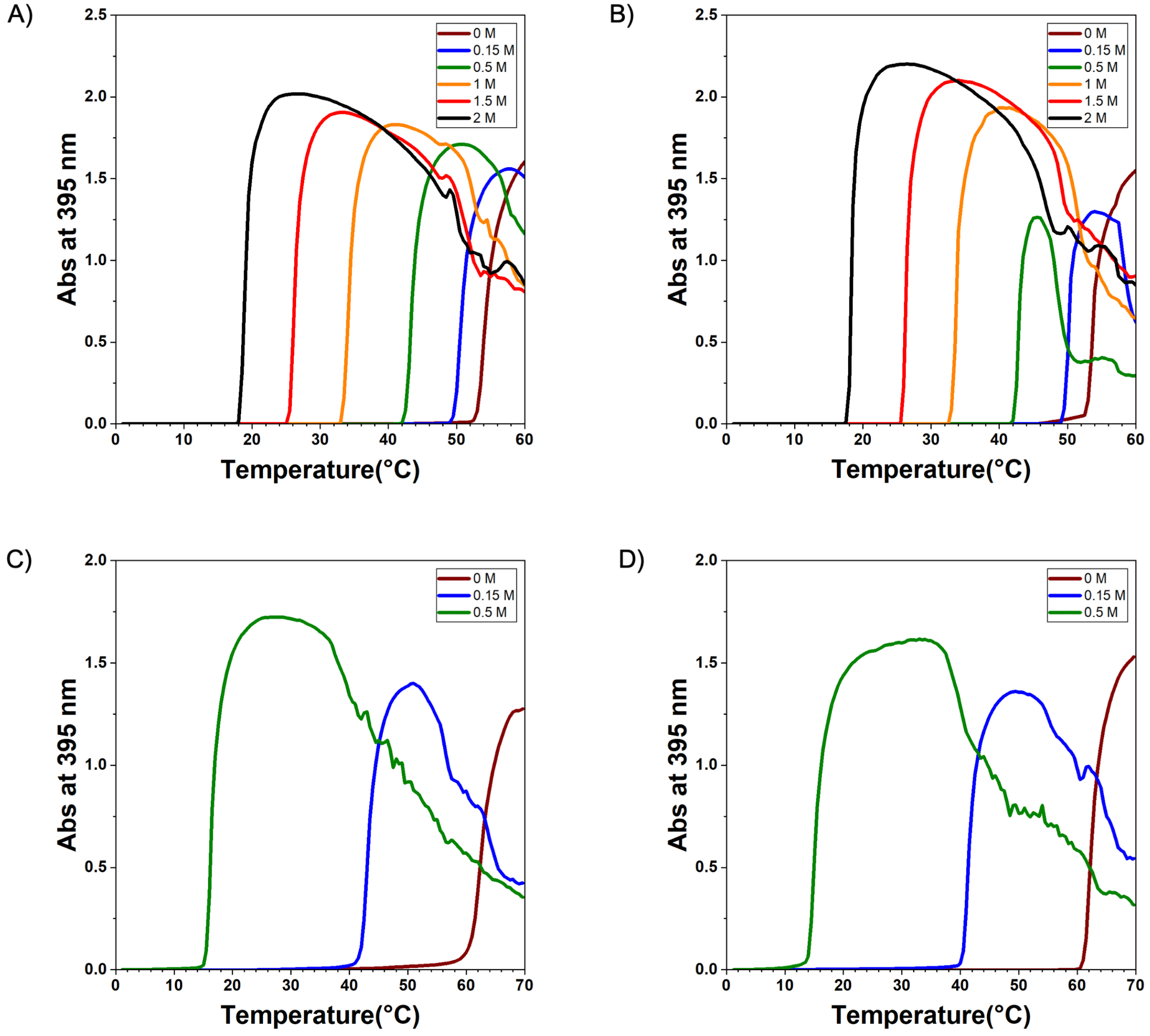


Figure S8: Experimental $T_t$ of ELP sequences. The $T_t$ diagram of (A) $V_{30}A_{30}$, (B) $A_{30}V_{30}$, (C) $V_{30}G_{30}$, and (D) $G_{30}V_{30}$. The turbidity was measured in the NaCl solution with different concentrations for $V_{30}A_{30}$ and $A_{30}V_{30}$ and in $(NH_4)_2SO_4$ solution with different concentrations for $V_{30}G_{30}$ and $G_{30}V_{30}$. The heat-up rate is 0.5 °C/min and absorbance data were collected every 0.5 °C.

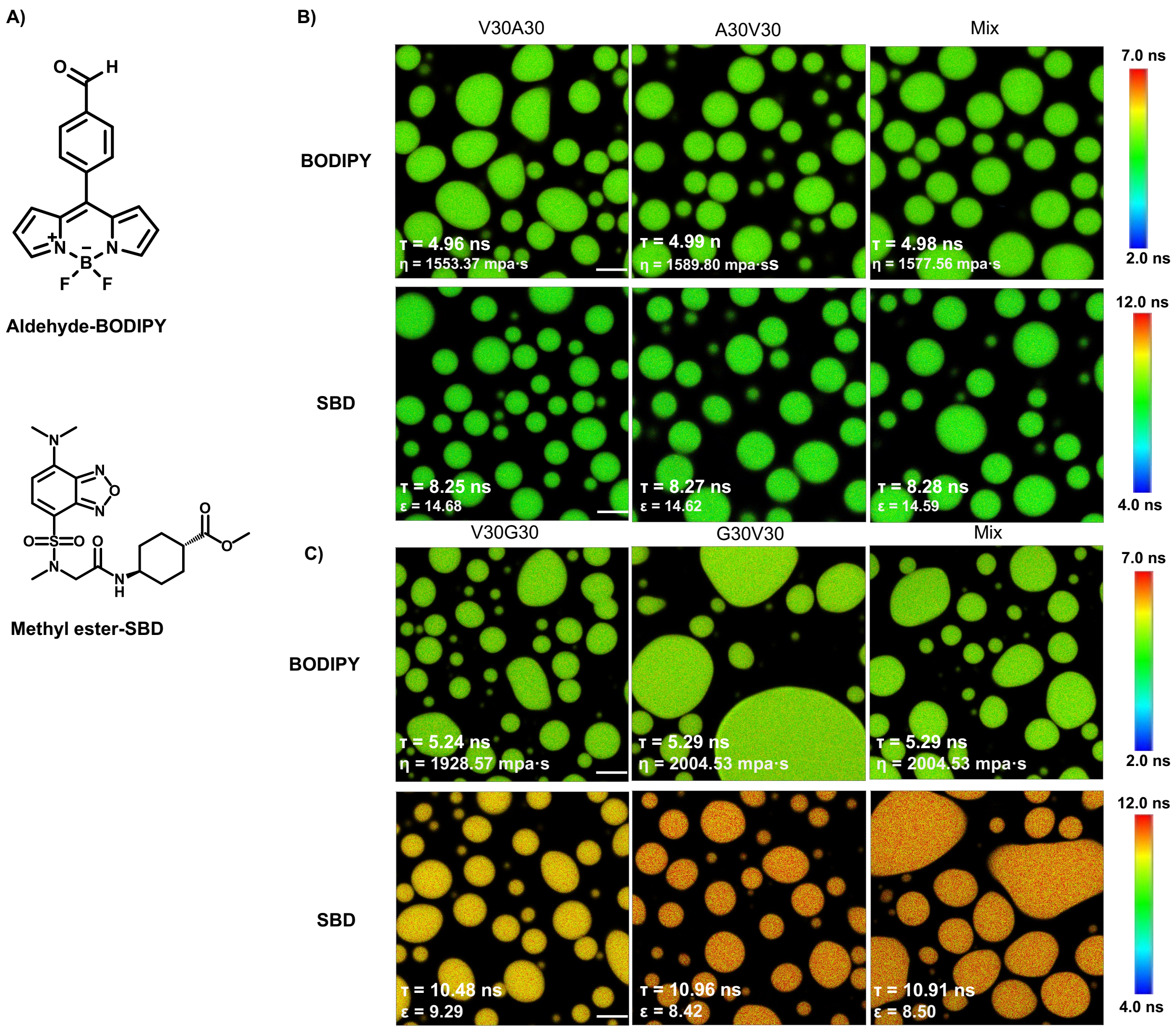


Figure S9: FLIM image of unlabeled ELP condensates. A) Chemical structure of free fluorophore, which can measure the physicochemical properties of condensates without labeling. B) Representative FLIM images of V30A30 and A30V30. The mix is the mixture of V30A30 (35 $\mu$M) and A30V30 (35 $\mu$M). Droplets were formed with a final concentration of 70 $\mu$M ELP in 2 M NaCl solution with 1 $\mu$M fluorophore. C) Representative FLIM images of V30G30 and G30V30. Droplets were formed with a final concentration of 70 $\mu$M ELP in 1.5 M (NH4)2SO4 solution with 1 $\mu$M fluorophore. The mix is the mixture of V30G30(35 $\mu$M) and G30V30 (35 $\mu$M). Scale bar, 5 $\mu$m. The fluorescence lifetime of each image is the average from three independent measurements.

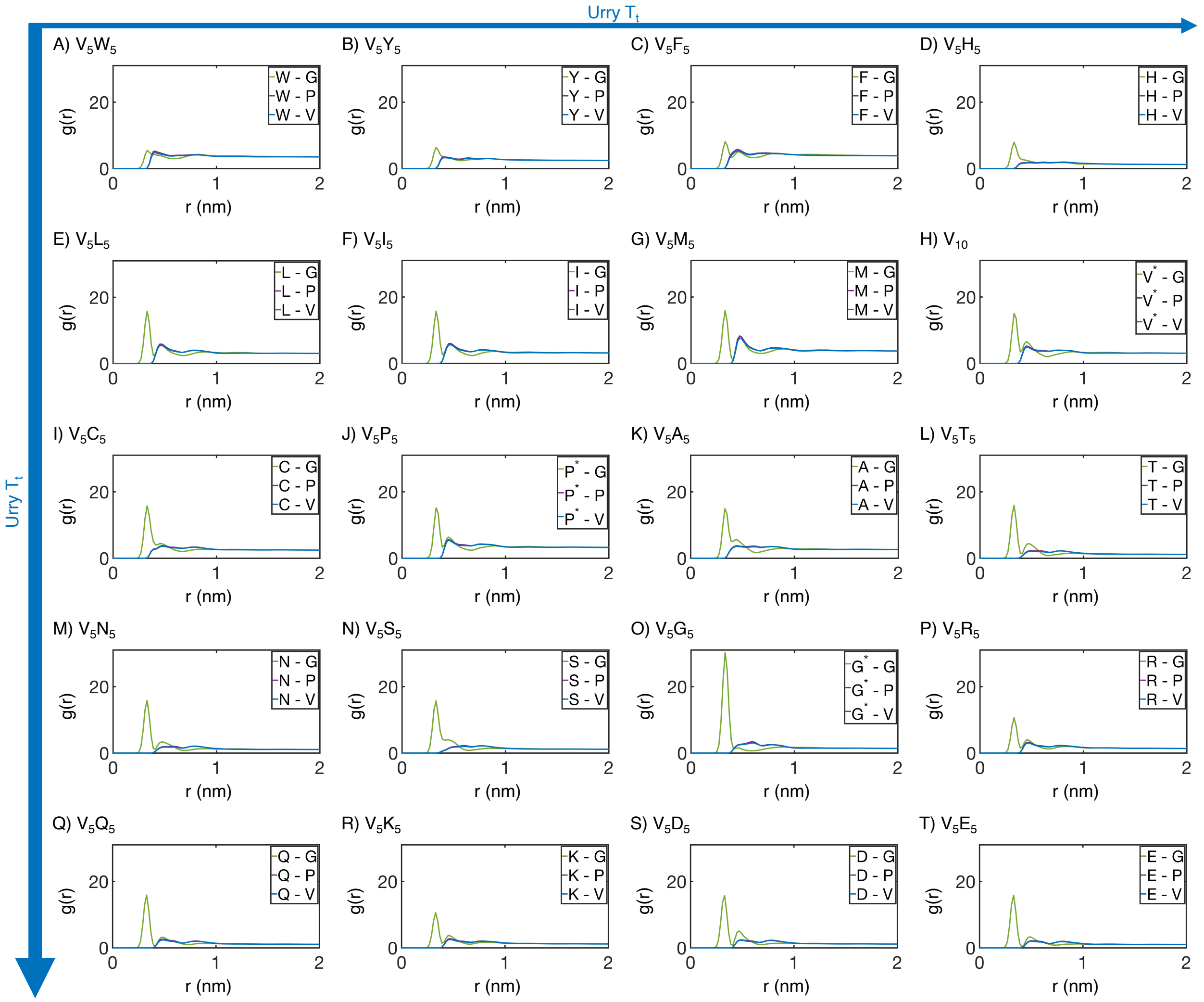


Figure S10: Residue specific radial distribution function $g(r)$ from MARTINI simulations. Distances from the guest amino acid (X) to those amino acids native to the ELP sequence were used to compute $g(r)$. Substituted amino acids that also appear in the native sequence are differentiated from their native counterparts using a star (*). Proteins are arranged by increasing $T_t$ (decreasing hydrophobicity) from Urry,[S22] with (A, $V_5W_5$) being the most hydrophobic and (T, $V_5E_5$) being the least hydrophobic.

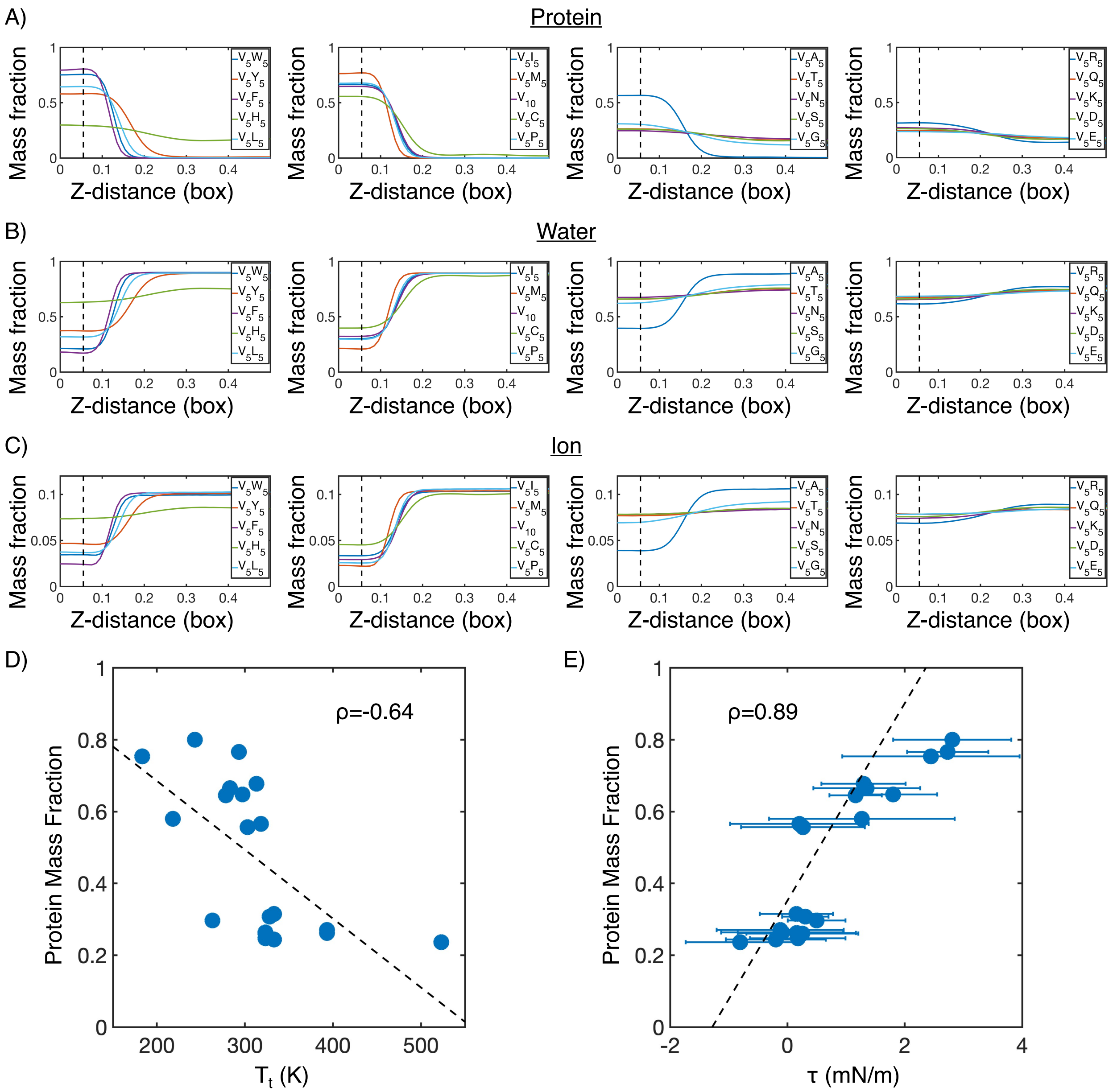


Figure S11: Mass fraction from MARTINI condensate simulations. (A) Protein mass fraction along the $Z$-distance from the condensate center defined using the largest cluster. The $Z-$axis is defined as the direction perpendicular to the condensate-water interface. The dashed line represents a $Z$-distance of 0.06 box lengths. Average values below this threshold are used for correlation analysis in parts D and E. (B) Water mass fraction along the $Z$-distance from the condensate center. (C) Ion mass fraction along the $Z$-distance from the condensate center. (D) Correlation between the average protein mass fraction and transition temperature ($T_{\mathrm{t}}$) from Urry.[S22] $\rho$ is the Pearson correlation coefficient between the two data sets, and the dashed diagonal line is the best fit line. Error bars represent standard deviations of the mean and are smaller than the symbols. (E) Correlation between the average protein mass fraction and the simulated surface tension ($\tau$). $\rho$ is the Pearson correlation coefficient between the two data sets, and the dashed diagonal line is the best fit line. Error bars represent standard deviations of the mean taken over 6 equally spaced box length intervals.

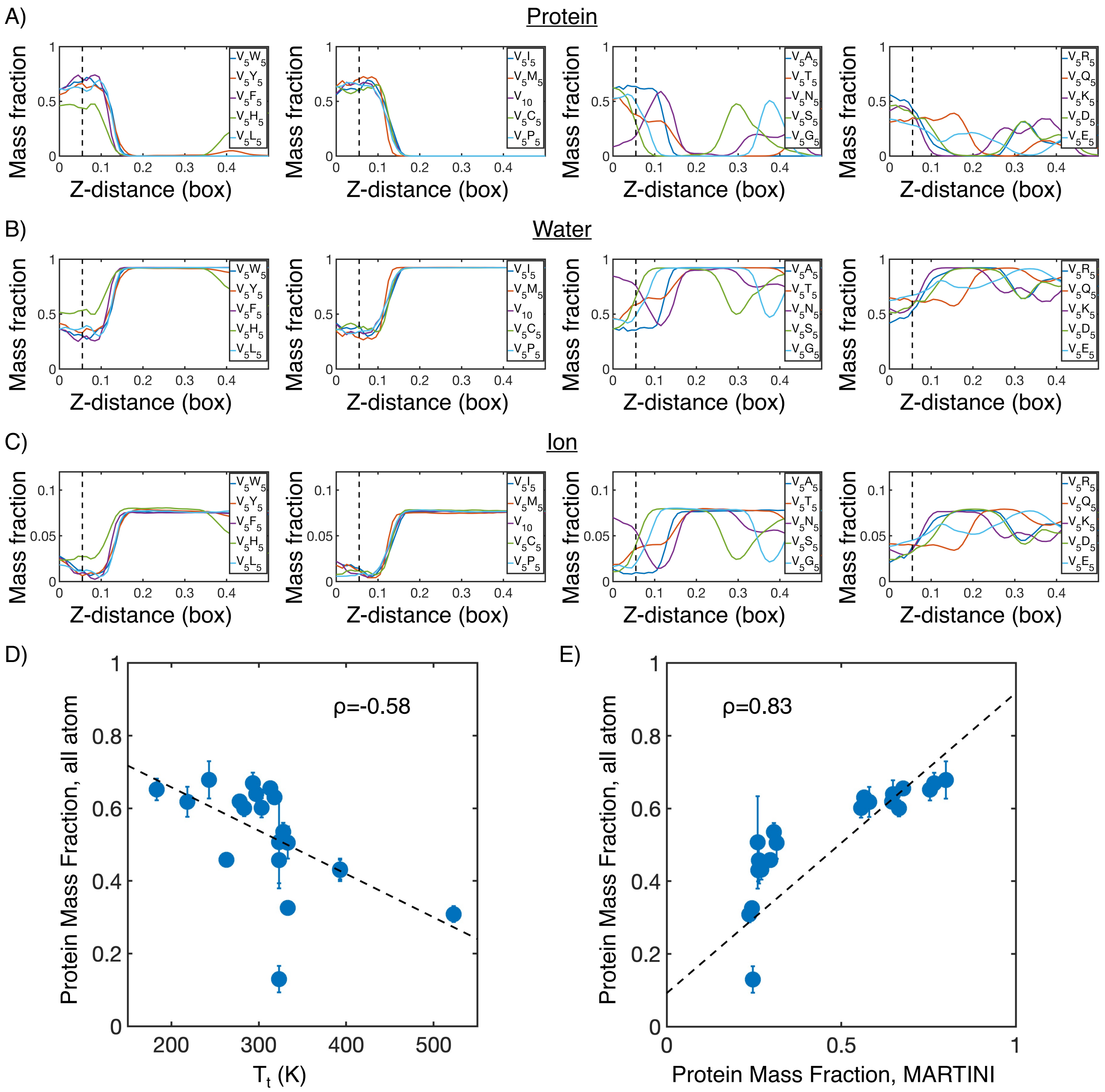


Figure S12: Mass fraction from atomistic condensate simulations. (A) Protein mass fraction along the $Z$-distance from the condensate center defined using the largest cluster. The $Z-$axis is defined as the direction perpendicular to the condensate-water interface. The dashed line represents a $Z$-distance of 0.06 box lengths. Average values below this threshold are used for correlation analysis in parts D and E. (B) Water mass fraction along the $Z$-distance from the condensate center. (C) Ion mass fraction along the $Z$-distance from the condensate center. (D) Correlation between the average protein mass fraction in all atom simulations and transition temperature ($T_{\mathrm{t}}$) from Urry.[S22] $\rho$ is the Pearson correlation coefficient between the two data sets, and the dashed diagonal line is the best fit line. Error bars represent standard deviations of the mean taken over 6 equally spaced box length intervals. (E) Correlation between the average protein mass fraction from all atom simulations and the average protein mass fraction from MARTINI simulations. $\rho$ is the Pearson correlation coefficient between the two data sets, and the dashed diagonal line is the best fit line. Error bars represent standard deviations of the mean taken over 6 equally spaced box length intervals.

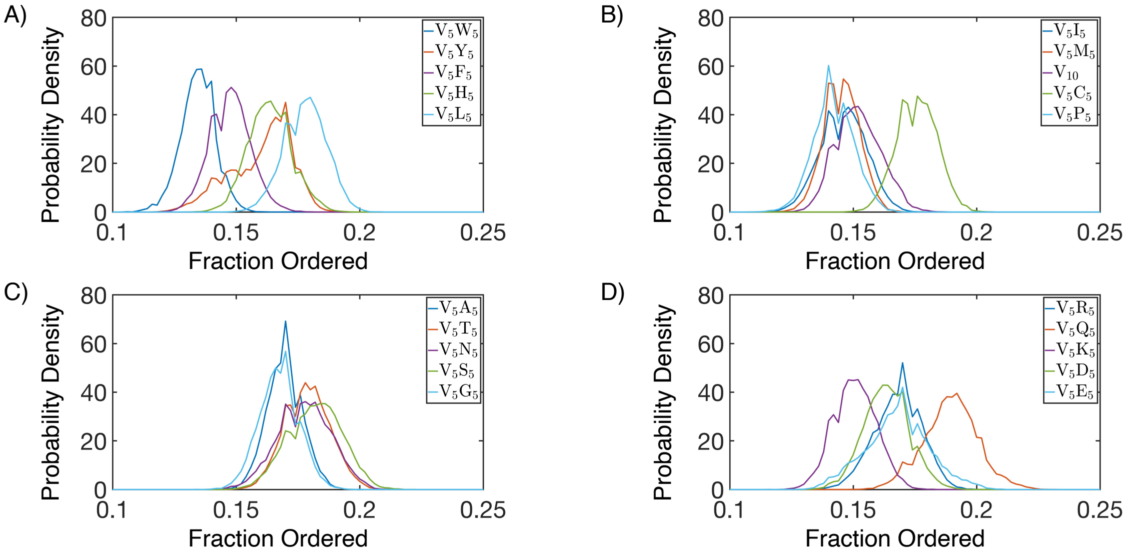


Figure S13: Secondary structure of simulated protein condensates. Secondary structure is defined according to the DSSP algorithm.[S32,S33] We consider $\alpha$-helices, $\beta$-sheets, $\beta$-bridges, and turns as ordered, and display the total fraction of all these secondary structure elements for each simulated condensate.

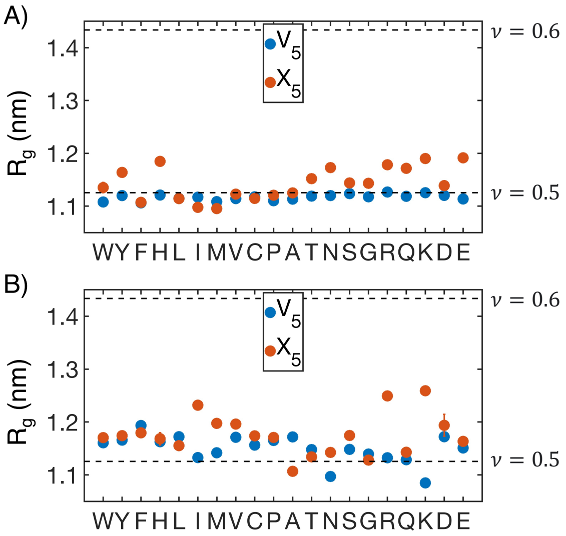


Figure S14: $R_g$ of individual peptide blocks from MARTINI (A) and atomistic (B) simulations. $V_5$ represents the valine block, while $X_5$ represents the guest block. Error bars represent standard deviations of the mean over 4 independent time windows. The dashed lines represent different values of polymer scaling exponents, with $\nu = 0.5$ corresponding to an ideal chain and $\nu = 0.6$ corresponding to a swollen polymer chain. Values previously suggested for IDPs were used in calculation of the scaling exponent.[S34]

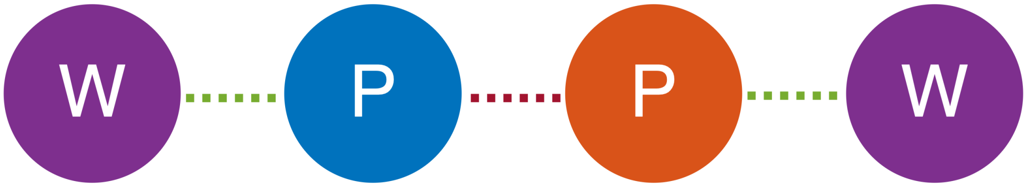


| | Protein-water | Protein-protein |
|---|---|---|
| Raw HB count | 2 | 1 |
| Per residue HB count | 1 ($\frac{n_{HB,P-W}}{n_{res,P}}$) | 1 ($\frac{2*n_{HB,P-P}}{n_{res,P}}$) |

Figure S15: Graphical illustration of number of hydrogen bonds per protein residue. The protein is depicted as blue and orange circles, and the water molecules are depicted as purple circles. Protein-protein and protein-water hydrogen bonds are drawn as red and green dashed lines respectively. The number of protein-water hydrogen bonds per protein residue is calculated by taking the raw number of protein-water hydrogen bonds ($n_{HB,P-W}$) and dividing by the number of residues in the protein ($n_{res,P}$). Meanwhile, the number of protein-protein hydrogen bonds per protein residue is calculated by doubling the number of protein-protein hydrogen bonds ($n_{HB,P-P}$) and dividing by the number of residues in the protein ($n_{res,P}$). The extra factor of 2 accounts for the fact that, in the case of protein-protein hydrogen bonds, both the donor and the acceptor must reside within the protein.

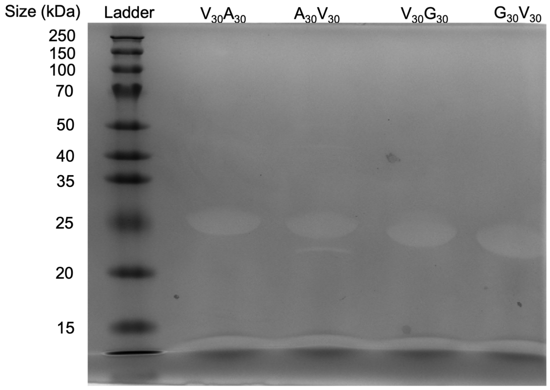


Figure S16: SDS-PAGE gel of ELP. The purity of the protein is > 95%.

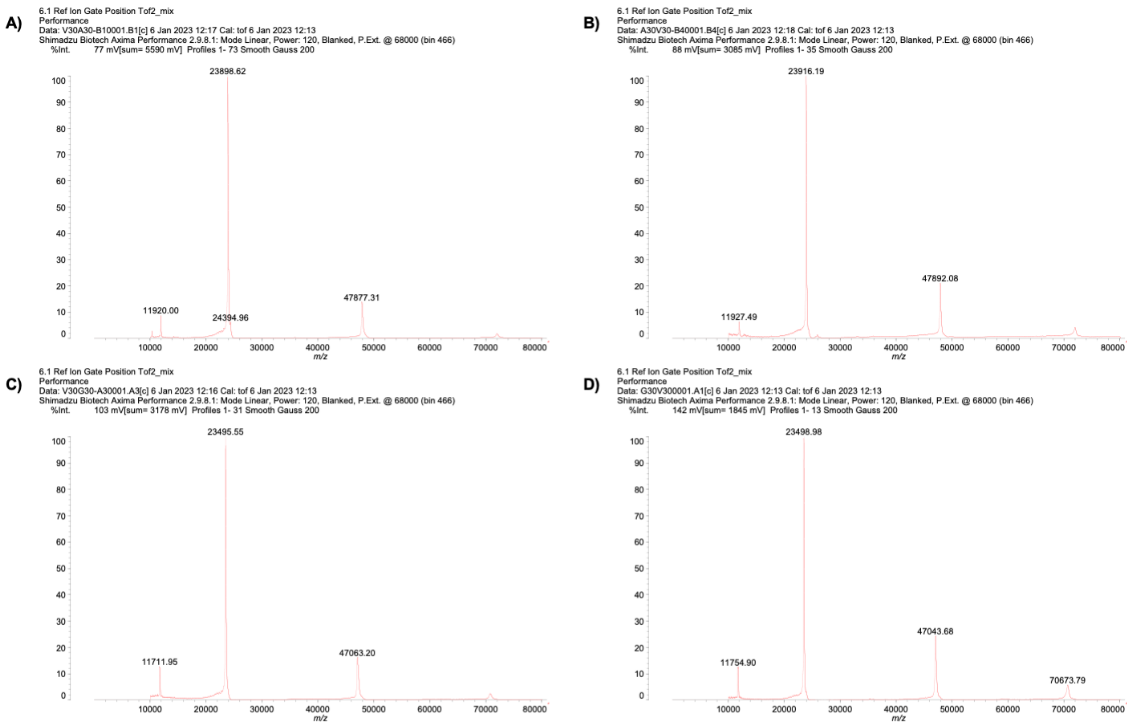


Figure S17: Mass spectrum of (A) $V_{30}A_{30}$, (B) $A_{30}V_{30}$, (C) $V_{30}G_{30}$, and (D) $G_{30}V_{30}$.

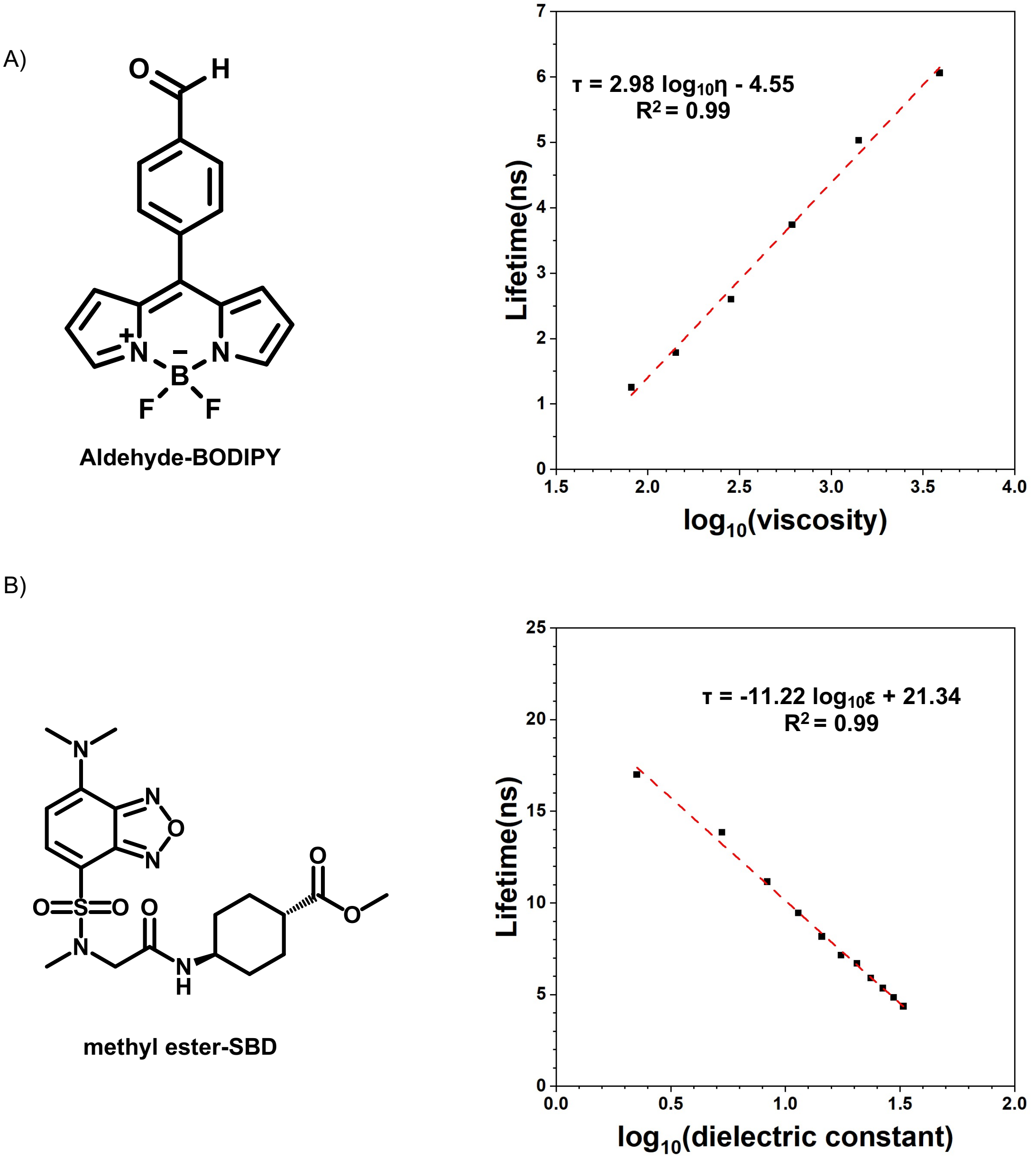


Figure S18: Calibration curves of fluorophores. (A) Lifetime-viscosity calibration curve is quantified in the glycerol at different temperatures (10 °C $\eta = 3900$ cP; 20 °C $\eta = 1412$ cP; 30 °C $\eta = 612$ cP; 40 °C $\eta = 284$ cP; 50 °C $\eta = 142$ cP; 60 °C $\eta = 81.3$ cP).[S35] The fluorophore concentration is 5 $\mu$M. (B) Lifetime-dielectric constant calibration curve is quantified in the MeOH - 1,4-dioxane mixture. $\epsilon_{mix} = \phi_1\epsilon_1 + \phi_2\epsilon_2$. $\epsilon_{MeOh} = 32.7$, $\epsilon_{1,4-dioxane} = 2.25$.

Table S1: Maximum allowed solvent accessible surface area (SASA) for each amino acid, taken from Tien et al.[S36]

| Amino Acid | maximum SASA (Å$^2$) |
|---|---|
| ALA | 129 |
| ARG | 274 |
| ASN | 195 |
| ASP | 193 |
| CYS | 167 |
| GLN | 225 |
| GLU | 223 |
| GLY | 104 |
| HIS | 224 |
| ILE | 197 |
| LEU | 201 |
| LYS | 236 |
| MET | 224 |
| PHE | 240 |
| PRO | 159 |
| SER | 155 |
| THR | 172 |
| TRP | 285 |
| TYR | 263 |
| VAL | 174 |

Table S2: Mass spectrum of purified ELP.

| | Mass (Da) | |
|---|---|---|
| | theoretical | MALDI-MS |
| $V_{30}A_{30}$ | 24097.04 | 23898.62 |
| $A_{30}V_{30}$ | 24097.04 | 23916.19 |
| $V_{30}G_{30}$ | 23676.23 | 23495.55 |
| $G_{30}V_{30}$ | 23676.23 | 23498.98 |

#### Experimental sequences

##### $\mathbf{V}_{30}\mathbf{A}_{30}$

M-$(GVGVP)_{30}$-$(GAGVP)_{30}$-GY

##### $\mathbf{A}_{30}\mathbf{V}_{30}$

M-$(GAGVP)_{30}$-$(GVGVP)_{30}$-GY

##### $\mathbf{V}_{30}\mathbf{G}_{30}$

M-$(GVGVP)_{30}$-$(GGGVP)_{30}$-GY

##### $\mathbf{G}_{30}\mathbf{V}_{30}$

M-$(GGGVP)_{30}$-$(GVGVP)_{30}$-GY

### References


(S1) Latham, A. P.; Zhang, B. Consistent Force Field Captures Homologue-Resolved HP1 Phase Separation. *J. Chem. Theory Comput.* **2021**, *17*, 3134–3144.

(S2) Souza, P. C. et al. Martini 3: a general purpose force field for coarse-grained molecular dynamics. *Nat. Methods* **2021**, *18*, 382–388.

(S3) Huang, J.; Rauscher, S.; Nawrocki, G.; Ran, T.; Feig, M.; de Groot, B. L.; Grubmüller, H.; MacKerell, A. D. CHARMM36m: an improved force field for folded and intrinsically disordered proteins. *Nat. Methods* **2016**, *14*, 71–73.

(S4) Berendsen, H. J.; van der Spoel, D.; van Drunen, R. GROMACS: A message-passing parallel molecular dynamics implementation. *Comput. Phys. Commun.* **1995**, *91*, 43–56.

(S5) Rauscher, S.; Pomès, R. The liquid structure of elastin. *eLife* **2017**, *6*, e26526.

(S6) Reichheld, S. E.; Muiznieks, L. D.; Keeley, F. W.; Sharpe, S. Direct observation of structure and dynamics during phase separation of an elastomeric protein. *Proc. Natl. Acad. Sci. U.S.A.* **2017**, *114*, E4408–E4415.

(S7) Mazur, J.; Jernigan, R. L. Distance-dependent dielectric constants and their application to double-helical DNA. *Biopolymers* **1991**, *31*, 1615–1629.

(S8) Mehler, E. L.; Solmajer, T. Electrostatic effects in proteins: Comparison of dielectric and charge models. *Protein Eng. Des. Sel.* **1991**, *4*, 903–910.

(S9) Case, D. et al. *Amber 2020*; University of California, San Francisco, 2020.

(S10) Dignon, G. L.; Zheng, W.; Kim, Y. C.; Best, R. B.; Mittal, J. Sequence determinants of protein phase behavior from a coarse-grained model. *PLoS Comput. Biol.* **2018**, *14*, 1–23.

(S11) Zhang, Y.; Feller, S. E.; Brooks, B. R.; Pastor, R. W. Computer simulation of liquid/liquid interfaces. I. Theory and application to octane/water. *J. Chem. Phys.* **1995**, *103*, 10252–10266.

(S12) Thomasen, F. E.; Pesce, F.; Roesgaard, M. A.; Tesei, G.; Lindorff-Larsen, K. Improving Martini 3 for Disordered and Multidomain Proteins. *J. Chem. Theory Comput.* **2022**, *18*, 2033–2041.

(S13) Benayad, Z.; Von Bülow, S.; Stelzl, L. S.; Hummer, G. Simulation of FUS Protein Condensates with an Adapted Coarse-Grained Model. *J. Chem. Theory Comput.* **2021**, *17*, 525–537.

(S14) Hassouneh, W.; Zhulina, E. B.; Chilkoti, A.; Rubinstein, M. Elastin-like Polypeptide Diblock Copolymers Self-Assemble into Weak Micelles. *Macromolecules* **2015**, *48*, 4183–4195.

(S15) Taylor, N.; Elbaum-Garfinkle, S.; Vaidya, N.; Zhang, H.; Stone, H. A.; Brangwynne, C. P. Biophysical characterization of organelle-based RNA/protein liquid phases using microfluidics. *Soft Matter* **2016**, *12*, 9142–9150.

(S16) Wassenaar, T. A.; Pluhackova, K.; Böckmann, R. A.; Marrink, S. J.; Tieleman, D. P. Going backward: A flexible geometric approach to reverse transformation from coarse grained to atomistic models. *J. Chem. Theory Comput.* **2014**, *10*, 676–690.

(S17) Baek, M. et al. Accurate prediction of protein structures and interactions using a three-track neural network. *Science* **2021**, *373*, 871–876.

(S18) Bonneau, R.; Tsai, J.; Ruczinski, I.; Chivian, D.; Rohl, C.; Strauss, C. E.; Baker, D. Rosetta in CASP4: Progress in ab initio protein structure prediction. *Proteins* **2001**, *45*, 119–126.

(S19) Dignon, G. L.; Zheng, W.; Best, R. B.; Kim, Y. C.; Mittal, J. Relation between single-molecule properties and phase behavior of intrinsically disordered proteins. *Proc. Natl. Acad. Sci. U.S.A.* **2018**, 201804177.

(S20) McDaniel, J. R.; MacKay, J. A.; Quiroz, F. G.; Chilkoti, A. Recursive directional ligation by plasmid reconstruction allows rapid and seamless cloning of oligomeric genes. *Biomacromolecules* **2010**, *11*, 944–952.

(S21) Meyer, D. E.; Chilkoti, A. Purification of recombinant proteins by fusion with thermally-responsive polypeptides. *Nat. Biotechnol.* **1999**, *17*, 1112–1115.

(S22) Urry, D. W. Physical chemistry of biological free energy transduction as demonstrated by elastic protein-based polymers. *J. Phys. Chem. B* **1997**, *101*, 11007–11028.

(S23) McDaniel, J. R.; Radford, D. C.; Chilkoti, A. A Unified Model for de Novo Design of Elastin-like Polypeptides with Tunable Inverse Transition Temperatures. *Biomacromolecules* **2013**, *14*, 2866–2872.

(S24) Meyer, D. E.; Chilkoti, A. Quantification of the Effects of Chain Length and Concentration on the Thermal Behavior of Elastin-like Polypeptides. *Biomacromolecules* **2004**, *5*, 846–851.

(S25) Flory, P. J. Thermodynamics of high polymer solutions. *J. Chem. Phys.* **1942**, *10*, 51.

(S26) Helfand, E.; Tagami, Y. Theory of the interface between immiscible polymers. *J. Chem. Phys.* **1972**, *56*, 3592.

(S27) Roe, R. J. Theory of the interface between polymers or polymer solutions. I. Two components system. *J. Chem. Phys.* **1975**, *62*, 490–499.

(S28) Rubinstein, M.; Colby, R. H. *Polymer Physics*; Oxford University Press, 2003.

(S29) Schauperl, M.; Podewitz, M.; Waldner, B. J.; Liedl, K. R. Enthalpic and Entropic Contributions to Hydrophobicity. *Journal of Chemical Theory and Computation* **2016**, *12*, 4600–4610.

(S30) Matsen, M. W.; Bates, F. S. Unifying weak- and strong-segregation block copolymer theories. *Macromolecules* **1996**, *29*, 1091–1098.

(S31) Swann, J. M.; Topham, P. D. Design and application of nanoscale actuators using block-copolymers. *Polymers* **2010**, *2*, 454–469.

(S32) Joosten, R. P.; Te Beek, T. A.; Krieger, E.; Hekkelman, M. L.; Hooft, R. W.; Schneider, R.; Sander, C.; Vriend, G. A series of PDB related databases for everyday needs. *Nucleic Acids Res.* **2015**, *43*, D364–D368.

(S33) Kabsch, W.; Sander, C. Dictionary of Protein Secondary Structure: Pattern Recognition of Hydrogen-Bonded and Geometrical. *Biopolymers* **1983**, *22*, 2577–2637.

(S34) Hofmann, H.; Soranno, A.; Borgia, A.; Gast, K.; Nettels, D.; Schuler, B. Polymer scaling laws of unfolded and intrinsically disordered proteins quantified with single-molecule spectroscopy. *Proc. Natl. Acad. Sci. U.S.A.* **2012**, *109*, 16155–16160.

(S35) Segur, J. B.; Oberstar, H. E. Viscosity of Glycerol and Its Aqueous Solutions. *Ind. Eng. Chem.* **1951**, *43*, 2117–2120.

(S36) Tien, M. Z.; Meyer, A. G.; Sydykova, D. K.; Spielman, S. J.; Wilke, C. O. Maximum allowed solvent accessibilites of residues in proteins. *PLoS ONE* **2013**, *8*.
